# Single residue substitution in protamine 1 disrupts sperm genome packaging and embryonic development in mice

**DOI:** 10.1101/2021.09.16.460631

**Authors:** Lindsay Moritz, Samantha B. Schon, Mashiat Rabbani, Yi Sheng, Devon F. Pendlebury, Ritvija Agrawal, Caleb Sultan, Kelsey Jorgensen, Xianing Zheng, Adam Diehl, Kaushik Ragunathan, Yueh-Chiang Hu, Jayakrishnan Nandakumar, Jun Z. Li, Alan P. Boyle, Kyle E. Orwig, Sy Redding, Saher Sue Hammoud

## Abstract

Conventional dogma presumes that protamine-mediated DNA compaction in sperm is achieved by passive electrostatics between DNA and the arginine-rich core of protamines. However, phylogenetic analysis reveals several non-arginine residues that are conserved within, but not across, species. The functional significance of these residues or post-translational modifications are poorly understood. Here, we investigated the functional role of K49, a rodent-specific lysine residue in mouse protamine 1 (P1) that is acetylated early in spermiogenesis and retained in sperm. *In vivo*, an alanine substitution (P1 K49A) results in ectopic histone retention, decreased sperm motility, decreased male fertility, and in zygotes, premature P1 removal from paternal chromatin. *In vitro*, the P1 K49A substitution decreases protamine-DNA binding and alters DNA compaction/decompaction kinetics. Hence, a single amino acid substitution outside the P1 arginine core is sufficient to profoundly alter protein function and developmental outcomes, suggesting that protamine non-arginine residues are essential to ensure reproductive fitness.

## Introduction

Spermatogenesis is a highly regulated differentiation process by which spermatogonial stem cells give rise to mature haploid spermatozoa throughout life. In its final stage, known as spermiogenesis, haploid round spermatids elongate to form sperm. During this morphological transformation, the global chromatin composition is completely altered, resulting in the near total replacement of histones by protamines. This transition from nucleosome-based to protamine-based chromatin in spermatids is a stepwise process. It begins with the exchange of canonical histones for testis-specific histone variants, such as spermatid-specific linker histone H1-like protein (HILS) (Yan et al., 2003), H2AL1/2 (Barral et al., 2017; Govin et al., 2007), testis-specific histone H2B (TH2B) (Montellier et al., 2013; Shinagawa et al., 2015), and histone H3T (Tachiwana et al., 2008, 2010; Ueda et al., 2017). Histone variants (both canonical and testis-specific) then acquire post-translational modifications (PTMs), notably hyperacetylation of H4 (Meistrich et al., 1992a; Shirakata et al., 2014) and ubiquitination of H2A/H2B (Lu et al., 2010), which initiates loosening of the chromatin structure to facilitate the incorporation of transition proteins 1 (TNP1) and 2 (TNP2), and subsequent replacement by protamines (Barral et al., 2017; Shirley et al., 2004a; Yu et al., 2000a).

Protamines are small, arginine-rich sperm-specific structural proteins that condense sperm chromatin (Pienta and Coffey, 1984; Ward and Coffey, 1991). Most mammals, including mice and humans, express two forms of protamine: protamine 1 (P1) and protamine 2 (P2). P1 is directly expressed in its mature form, while P2 is expressed as a longer precursor (pro P2) that is initially deposited onto DNA and subsequently processed by a series of selective proteolytic cleavages to produce its mature form (P2) (Green et al., 1994; Yelick et al., 1987). Together, P1 and P2 wrap 90-95% of the mammalian sperm genome (Ward and Coffey, 1991; Wykes and Krawetz, 2003). Numerous studies in both mice and humans have demonstrated that maintenance of a species-specific ratio of P1:P2 (1:1 in humans, 1:2 in mouse) is necessary for fertility, and that alterations in this ratio correlate with decreased fertility and poor embryonic development (Aoki et al., 2006; de Mateo et al., 2009; Zatecka et al., 2014). Furthermore, from knockout and haploinsufficiency studies, we know that P1 and P2 are essential for sperm chromatin packaging and fertility (Cho et al., 2001, 2003; Schneider et al., 2016; Takeda et al., 2016a).

Although protamines’ role in packaging the majority of the sperm genome and requirement for fertility are widely known, the regulation of mammalian protamine-induced DNA condensation and subsequent decondensation in the zygote is not well understood. Much of our understanding of protamine-DNA dynamics arises from early *in vitro* studies utilizing either salmon or bull (domestic cattle, *Bos taurus*) sperm protamine, both of which express only one form of protamine (Balhorn et al., 2000; Bench et al., 1996; Brewer, 1999; Brewer et al., 2003; Prieto et al., 1997a). However, most mammalian genomes encode multiple protamine proteins (P1, P2, and/or P3) that may engage in complex inter and intramolecular interactions between different protamine forms, which cannot be captured or monitored in species which encode for a single protamine protein (Balhorn et al., 2000; Brewer et al., 1999; Prieto et al., 1997b) or species that lack cysteine residues (like Salmon). Hence, our understanding of a complex, multi-protamine packaging system is limited and based on the assumption that the biochemical and biophysical properties of protamines are conserved, despite striking differences in amino acid sequence and composition across animal species (Krawetz and Dixon, 1988; Lewis et al., 2003; Wyckoff et al., 2000). Consequently, the framework based on our current knowledge is unlikely to accurately describe functional differences and/or kinetics of mammalian protamines or systems that employ a dual protamine (P1 and P2) packaging system.

In somatic cells, histone proteins package the DNA, and these proteins acquire various PTMs which impact histone-DNA interaction strength and chromatin and transcriptional states (Brehove et al., 2015; Kiefer et al., 2008; Shogren-Knaak, 2006). In sperm, a similar series of modifications have been reported for mouse and human protamines, but given the basic nature of protamine proteins and apparent evolutionary selection for high arginine content, most studies of protamine-DNA interactions centered on arginine residues, limiting our understanding of functional roles for other residues or protamine PTMs (Queralt et al., 1995; Rooney et al., 2000; Torgerson et al., 2002). However, a few studies that examined protamine phosphorylation and dephosphorylation have suggested that protamine phosphorylation during spermiogenesis is important for modulating protamine-DNA dynamics and maximizing chromatin compaction (Green et al., 1994; Itoh et al., 2019; Pirhonen et al., 1994; Seligman et al., 2004). More recently, Guo. et. al. reported that several serine residues in P1 acquired phosphorylation during early embryogenesis and these modifications were required to weaken protamine-DNA interactions to allow male pronuclear remodeling and protamine-to-histone exchange—further supporting a model where electrostatic interactions are the primary mode of regulation of sperm chromatin organization (Gou et al., 2020).

To examine whether protamine-DNA interactions can solely be explained by simple electrostatics or if the protamine sequence itself and/or the dynamics of its modifications are instructive for proper chromatin packaging in sperm and unpackaging in the early embryo, we performed mass spectrometry and phylogenetic analysis of P1 sequences. We find that the sites of protamine PTMs are conserved within species, but not across, suggesting the possibility of lineage specific function. To dissect this phenomenon genetically, we chose to focus on P1 lysine 49—a residue that is conserved across the rodent lineage and we find to be acetylated in mature mouse sperm. Specifically, we report that P1 K49 acetylation is acquired in early elongating spermatids (stage IX-XI) and persists in mature sperm. The substitution of K49 with alanine (A) results in severe male subfertility in mice. Biochemical analysis of mutant sperm nuclei reveals alterations in sperm chromatin composition and histone eviction. K49A mutant embryos prematurely dismiss P1 from paternal chromatin and many of these embryos arrest at the 1-cell and blastocyst stages. *In vitro*, bulk and single molecule assays reveal that the K49A mutant protein has significantly lower affinity for DNA, slower rates of DNA condensation, and accelerated de-condensation— consistent with premature dismissal of P1 in embryos. All together, these findings establish an indispensable role for P1 amino acids outside of a general electrostatic model, highlighting a more complex role for protamine protein sequence in governing protamine-DNA genome packaging and embryonic development.

## Results

### Post-translational modifications on P1 are lineage specific

Previous top-down and bottom-up mass spectrometry studies identified several P1 and P2 PTMs in both human and mouse mature sperm; however, their function (aside from observations of P1 phosphorylation in the early embryo and P2 phosphorylation/dephosphorylation during spermatogenesis) remains unknown (Brunner et al., 2014; Gou et al., 2020; Itoh et al., 2019; Soler-Ventura et al., 2020). We were intrigued by these data, but because sperm are transcriptionally quiescent, these modifications cannot be involved in germ cell transcriptional regulation. Therefore, we set out to investigate possible alternative functions of protamine PTMs. First, we used mass spectrometry to independently validate that these modifications are present in mature mouse sperm. Through this analysis, we confirmed previously identified P1 modifications, such as phosphorylation at serine (S) 9, S43, and threonine (T) 45 and acetylation at lysine (K) 50, but we also identified additional modifications, such as acetylation of P1 at K49 (P1 K49ac) (**Supplemental Table 1**, summarized in **Figure 1A**). To gain a deeper understanding of sequence conservation of PTM-bearing sites, we constructed a phylogenetic tree for species across the orders Rodentia, Primate and Artiodactyla (hoofed animals, including *Bos taurus*) using maximum likelihood inferred from P1 protein sequences (**Figure 1B**). We found that the P1 S9 position is highly conserved, and its phosphorylation is well-established in both mouse and human, likely reflecting that both the amino acid position and modification serve a necessary function across species (Brunner et al., 2014; Chira et al., 1993). In contrast, several C-terminal modified residues (S43, T45, K49, K50) are all highly conserved within the mouse lineage but are all largely occupied by alternative residues in more distant species (**Figure 1B**). Given that the K49 residue is conserved across rodents but not primates, we developed a host of molecular and genetic reagents to begin to dissect the role of non-arginine residues, and specifically explore the functional role of K49 in the mouse germline.

**Figure 1:**
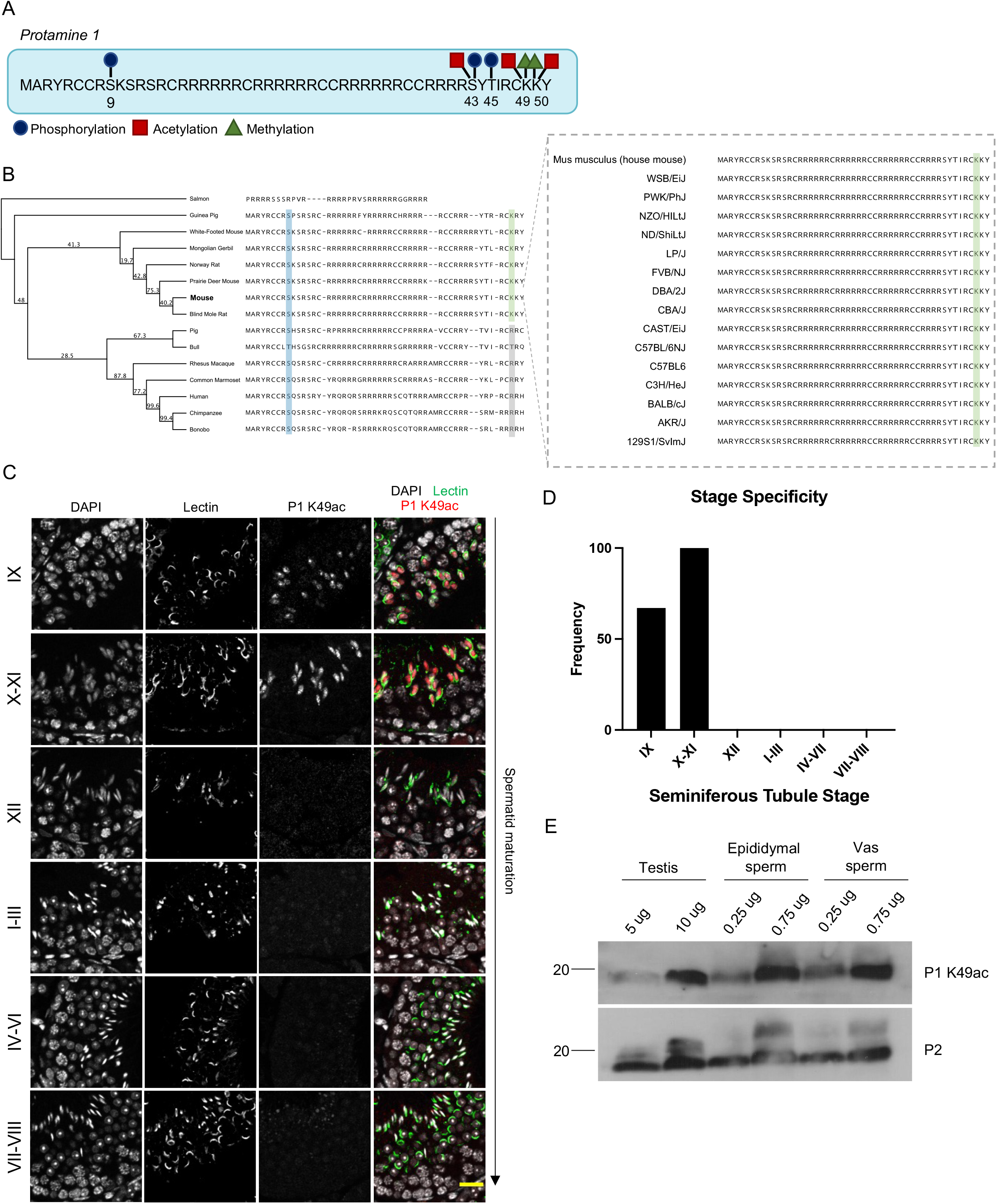
P1 lysine 49 acetylation is acquired in the testis in a stage-specific manner and is present in mouse sperm. **(A)** Schematic representation of modifications identified on mouse P1 using bottom-up mass spectrometry. **(B)** Phylogenetic tree constructed using maximum likelihood inferred from P1 protein sequences for species across the orders Rodentia, Primate, and Artiodactyla, using the WAG substitution strategy. Bootstrap support with 1,000 replicates is shown for each node, with values >95 indicating strong support. S9 is highlighted across species in blue and K49 is highlighted across rodents in green and in gray across more distant species that occupy alternative residues at this site. **(C)** Immunofluorescence staining of P1 K49ac in adult testes cross sections at various seminiferous tubule stages using PNA-Lectin as the acrosomal marker. Representative images from n=4 mice. Scale bar: 20 μm. **(D)** Quantification of P1 K49ac stage specificity across all stages, highlighting specificity to stage IX-XI tubules. A total of n=438 tubules were counted across all stages from a total of n=4 mice. **(E)** Western blot analysis of P1 K49ac from elongating spermatid-enriched testes lysate, mature sperm from the epididymis, and mature sperm from the vas deferens highlights the persistence of the acetylation mark into mature sperm.

### P1 K49 acetylation is acquired in the testis in a stage-specific manner and persists in mature sperm

To detect the presence of K49 acetylation (K49ac) in sperm, we generated a polyclonal antibody against P1 K49ac. In immunoblots of acid extracted protein from mature sperm, we found that our antibody detected a distinct band. This band was lost when outcompeted by an acetylated P1 K49 peptide, but not when we used a nonspecific peptide from an unrelated protein, or a non-acetylated P1 peptide, thus confirming both the presence of K49ac *in vivo* and the specificity of our antibody (**Figure S1A**). To precisely define at which stage or stages of the seminiferous tubule cycle P1 K49ac is established, we co-stained testes cross-sections using our custom antibody combined with the acrosomal marker PNA-Lectin. We found that specific signal was initiated in stage IX (containing early elongating spermatids) and peaked at stages X-XI (100% of tubules) but then diminished in stages XII-VIII (late-stage spermatids, **Figure 1C,D, S1B**). Although the fluorescent signal for P1 K49ac is diminished in later stages of spermatid maturation, the modification remained detectable in elongating spermatid-enriched lysates from the testis, as well as in epididymal and vas deferens sperm by immunoblotting (**Figure 1E**). These data point to two possible interpretations: loss of signal may be the result of high compaction of spermatids or low abundance below the immunofluorescence detection limit for our antibody.

### Substitution of P1 K49 for alanine results in sperm motility defects, abnormal sperm morphology, and subfertility

To investigate the functional significance of P1 K49ac *in vivo*, we used CRISPR/Cas9 to generate a lysine to alanine mutant mouse (K49A). We then used Sanger sequencing to confirm the presence of the target mutation (**Figure 2A**) and the absence of potential off-target genetic modifications (**Figure S2A**). Overall, P1^K49A/+^ or P1^K49A/K49A^ mice appeared phenotypically normal; we observed no significant differences in testes/body weight ratio and all germ cell populations were detected (**Figure S2B**,**C**). However, while overall sperm counts were normal in P1^K49A/K49A^ mice, progressive sperm motility (the ability of sperm to swim forward) was severely impaired (**Figure 2B,C**) and various sperm structural abnormalities were noted including coiled midpieces, bent back heads, and abnormal head morphology (**Figure 2D,E**). Furthermore, P1^K49A/K49A^ males were severely subfertile (**Figure 2F**). To ensure that this phenotype is not caused by loss of the P1 protein itself, we stained both P1^+/+^ and P1^K49A/K49A^ testis cross-sections using a custom P1 antibody (**Figure S2D, specificity test in Figure S2E**). We found that P1 is detectable in both cases, and that P1 protein levels are comparable between genotypes (**Figure S2F, 3A**), suggesting that the phenotype is not simply due to loss of P1 expression.

**Figure 2:**
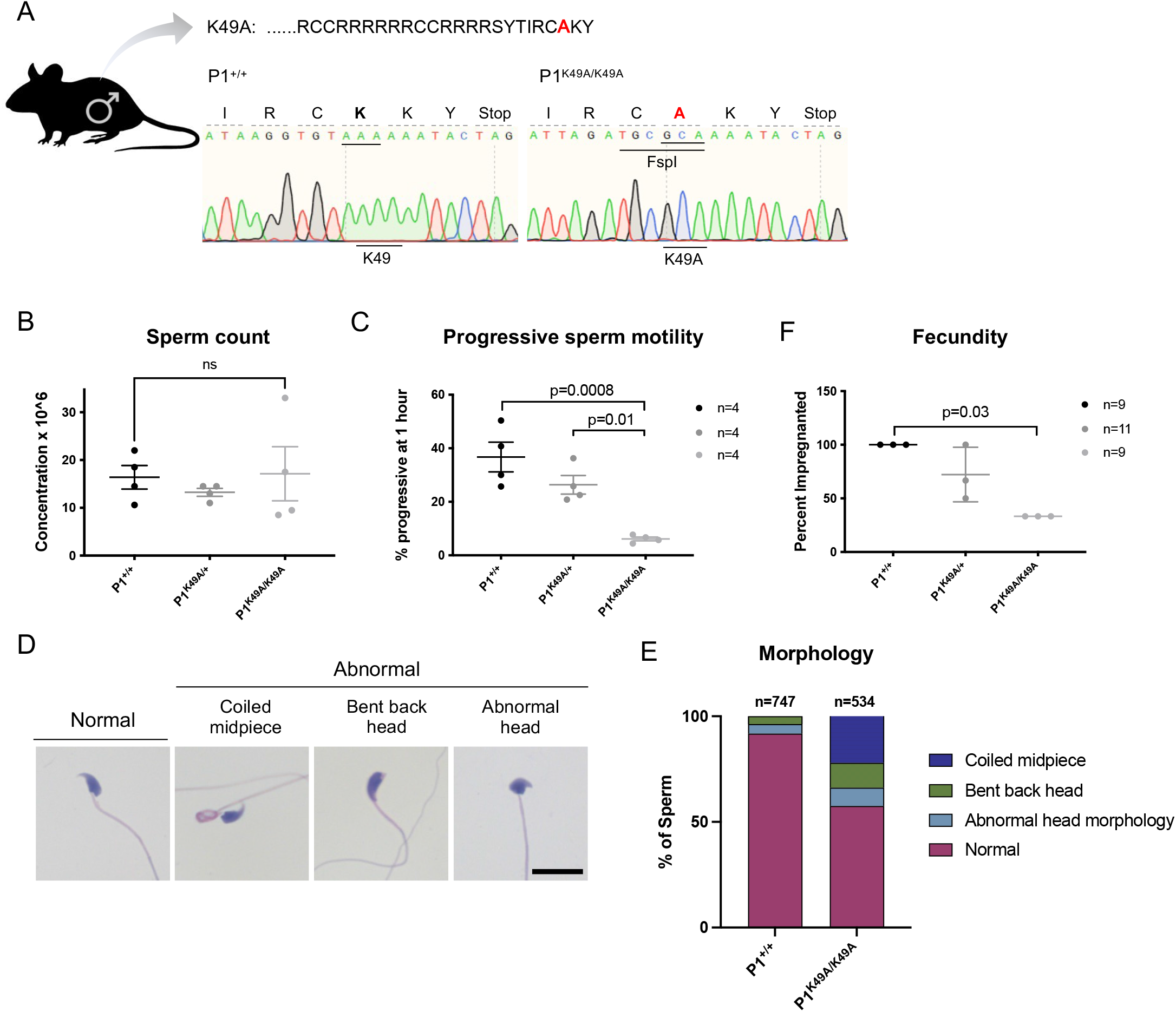
P1 K49A substitution results in sperm motility defects and subfertility. **(A)** Schematic of modification made to the mouse P1 sequence and corresponding Sanger sequencing traces illustrating successful mutation of K49 to alanine. Note that several synonymous mutations were incorporated into the donor DNA to prevent recutting of the target allele, which also introduced an FspI site that was utilized for genotyping. **(B-C)** Total epididymal sperm count (**B**, n=4 for each genotype) and epididymal sperm progressive motility after 1 hour of incubation at 37 °C (**C**, n=4 for each genotype.) Statistical tests were performed using a one-way ANOVA and adjusted for multiple comparisons. **(D)** Representative Hematoxylin and eosin-stained mature sperm from a P1^K49A/K49A^ adult male, highlighting observed major abnormalities observed. Scale bar: 20 μm. **(E)** Quantification of major abnormalities observed in P1^+/+^ and P1^K49A/K49A^ mature sperm. Sperm was assessed from n=3 P1^+/+^males and n=3 P1^K49A/K49A^ males. **(F)** Fertility assessment of 3 adult males per genotype as measured by percent of females impregnated (fecundity). Statistical test was performed using a Kruskal-Wallis test, p=0.03.

### P1 K49A mutants progress through key chromatin intermediate stages of the histone-to-protamine exchange, yet have abnormal histone retention

Given the abnormal sperm motility and morphology, we next analyzed the effects of the K49A substitution on the histone-to-protamine exchange and mature sperm chromatin composition. When we compared protamine levels and ratios in a fixed number of P1^+/+^, P1^K49A/+^, and P1^K49A/K49A^ sperm, we found that ratios in P1^K49A/K49A^ sperm were significantly shifted from the expected 1:2 P1:P2 ratio, to a ratio closer to ∼1:1. This decrease in P1:P2 ratio is not caused by a change in P1 levels, but is instead the result of accumulation of unprocessed P2 (pro P2, **Figure 3A**). Despite the defects in P2 processing, the total level of P2 (processed and unprocessed) remained unchanged, and the corrected ratio using P1: total P2 was ∼1:2.4. These results collectively suggest that the P1 K49A substitution does not affect P1 or P2 expression or overall protamine levels in sperm, but does affect the amount of processed P2 in sperm. However, we found that P1^K49A/K49A^ sperm had ∼3.5 fold higher levels of histones retained than P1^+/+^ (**Figure 3B,C**), suggesting that the P1 K49A protein disrupts overall histone eviction.

**Figure 3:**
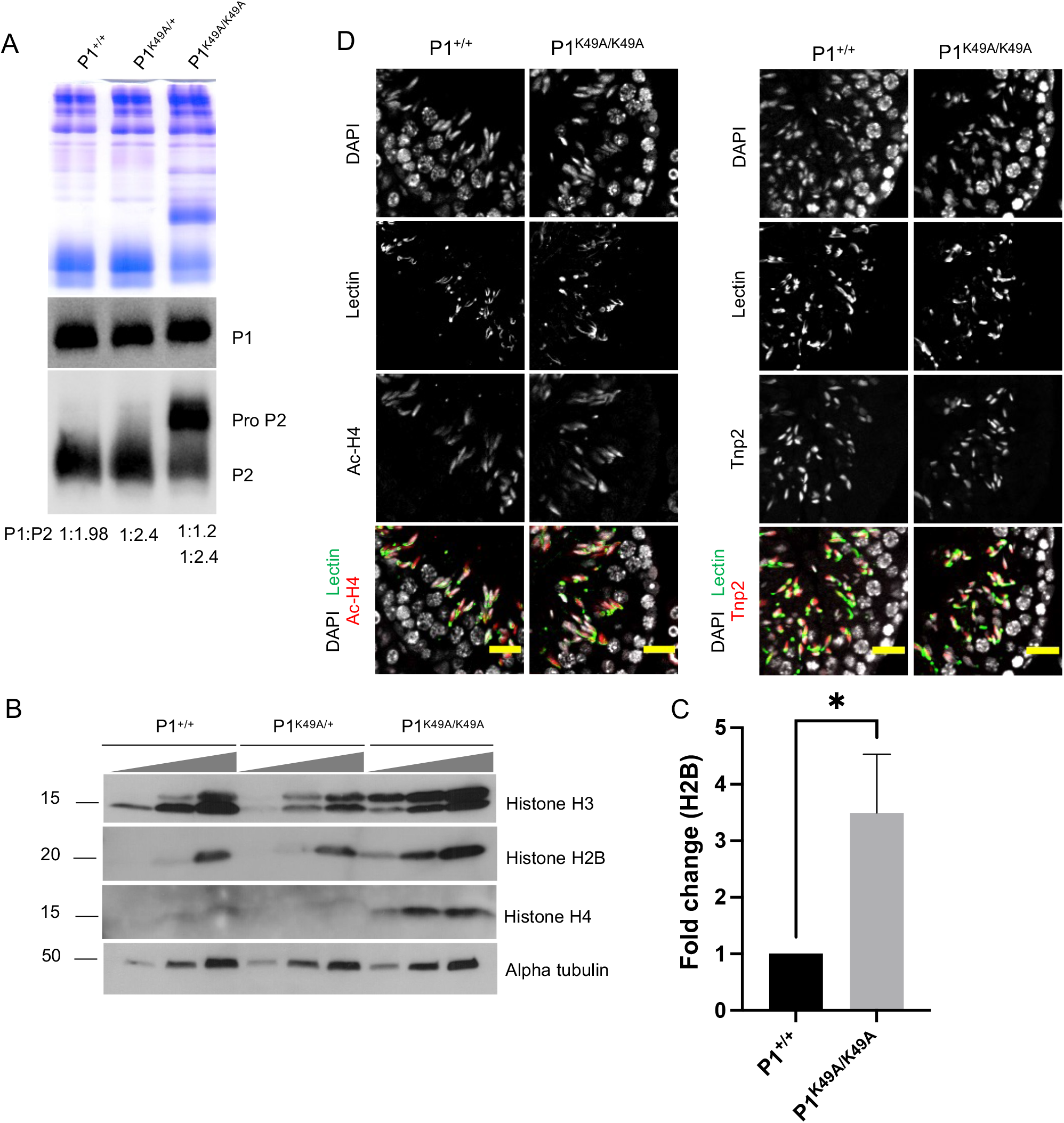
P1 K49A substitution alters sperm chromatin composition. **(A)** Acid urea gel electrophoresis of sperm basic proteins reveals a dramatic shift in P1:P2 ratio in P1^K49A/K49A^ males by Coomassie blue staining (top). Immunoblotting of the gel reveals no difference in P1 level but an accumulation of pro P2 (bottom panels). P1:P2 ratios as quantified in ImageJ are displayed below the immunoblot. **(B)** Immunoblotting of sperm protein extracts reveals an abnormal retention of histones in P1^K49A/K49A^ sperm. Blots were loaded by total input sperm number. Exact sperm numbers for the various antibodies provided in Methods section. **(C)** Quantification of immunoblots showing fold change of histone retention in P1^K49A/K49A^ males. Histone H2B was used as a representative protein for quantification and quantification was performed from n=4 technical replicates; sperm was pooled from n=2 mice. Statistical test was performed using a Mann-Whitney test, p=0.0286. **(D)** Immunofluorescence staining of adult P1^+/+^ or P1^K49A/K49A^ testes cross sections for ac-H4 (left panels) or Tnp2 (right panels). Scale bar: 20 μm.

As H4 hyperacetylation is indispensable for histone-to-protamine exchange (Dong et al., 2017; Ketchum et al., 2018; Luense et al., 2019; Meistrich et al., 1992b; Shiota et al., 2018), we next asked whether such initial triggering events occurred normally in P1^K49A/K49A^ mice. To this end, we analyzed P1^+/+^ and P1^K49A/K49A^ testes using an anti-H4 tetra-acetyl (referred to as ac-H4) antibody but observed no significant difference in ac-H4 levels by immunostaining (**Figure 3D, left panels**) or immunoblotting of testes lysates (**Figure S3A**). Next, we investigated transition proteins (TNP1 and TNP2), well known chromatin intermediate components of the histone-to-protamine exchange. Specifically, loss of TNP1 and TNP2 perturbs sperm morphology, chromatin composition, and final chromatin packaging—similar to our observations in P1^K49A/K49A^ mice (Shirley et al., 2004a; Yu et al., 2000a). When we analyzed TNP1 and TNP2 proteins in both testis cross-sections by immunostaining and testes lysates by immunoblotting, we did not observe any overt differences when comparing P1^+/+^ and P1^K49A/K49A^ mice (**Figure 3D, S3A**,**C**). Taken together, the P1^K49A/K49A^ mutants appear to progress normally through several key intermediate processes, yet the ultimate chromatin packaging is strikingly abnormal. These observations raise the question of whether acetylation of K49 itself may be required in the remodeling process, or whether other intermediate histone variants may be improperly loaded.

We next asked whether the retained histones in P1^K49A/K49A^ sperm are selectively enriched for specific histone PTMs which would indicate possible regional or programmatic alterations in histone eviction. To answer this question, we performed histone PTM immunoblots of protein extracts from increasing numbers of P1^+/+^, P1^K49A/+^ and P1^K49A/K49A^ sperm, and probed for a series of both activating and repressive modifications including ac-H4, H3K27ac, H3K9me3, and H3K27me3, and H4K20me3 (**Figure S3B**). When comparing across genotypes, it was evident that all PTMs analyzed appeared to be globally increased in the P1^K49A/K49A^ mutant, except for ac-H4, which was consistently lower in multiple biological replicates (**Figure S3B, data not shown**). In conclusion, the P1 K49A substitution does not compromise P1 protein stability, but rather the mutation causes functional changes to sperm chromatin, likely compromising the ability of protamines to compete against stably bound nucleosomes.

### P1 K49A substitution results in decreased blastocyst formation and accelerates P1 dismissal from paternal chromatin

Given the severely reduced motility observed in the sperm of P1^K49A/K49A^ males and severe subfertility, we next performed intracytoplasmic sperm injections (ICSI) to examine early developmental consequences of the P1 K49A substitution. In agreement with our natural mating data, blastocyst formation was significantly impaired when we injected P1^K49A/K49A^ sperm (13.5% vs. 48.0% using P1^+/+^ sperm, **Figure 4A**). When assessing embryo development and survival every ∼24 hours, we noticed two significant blocks in development in embryos derived from P1^K49A/K49A^ sperm: the first at the 1-cell to 2-cell transition, and, surprisingly, the other from morula to blastocyst stages (**Figure 4A**). This later block is striking and suggestive of abnormal transcriptional or epigenetic landscapes caused by the P1 K49A substitution.

**Figure 4:**
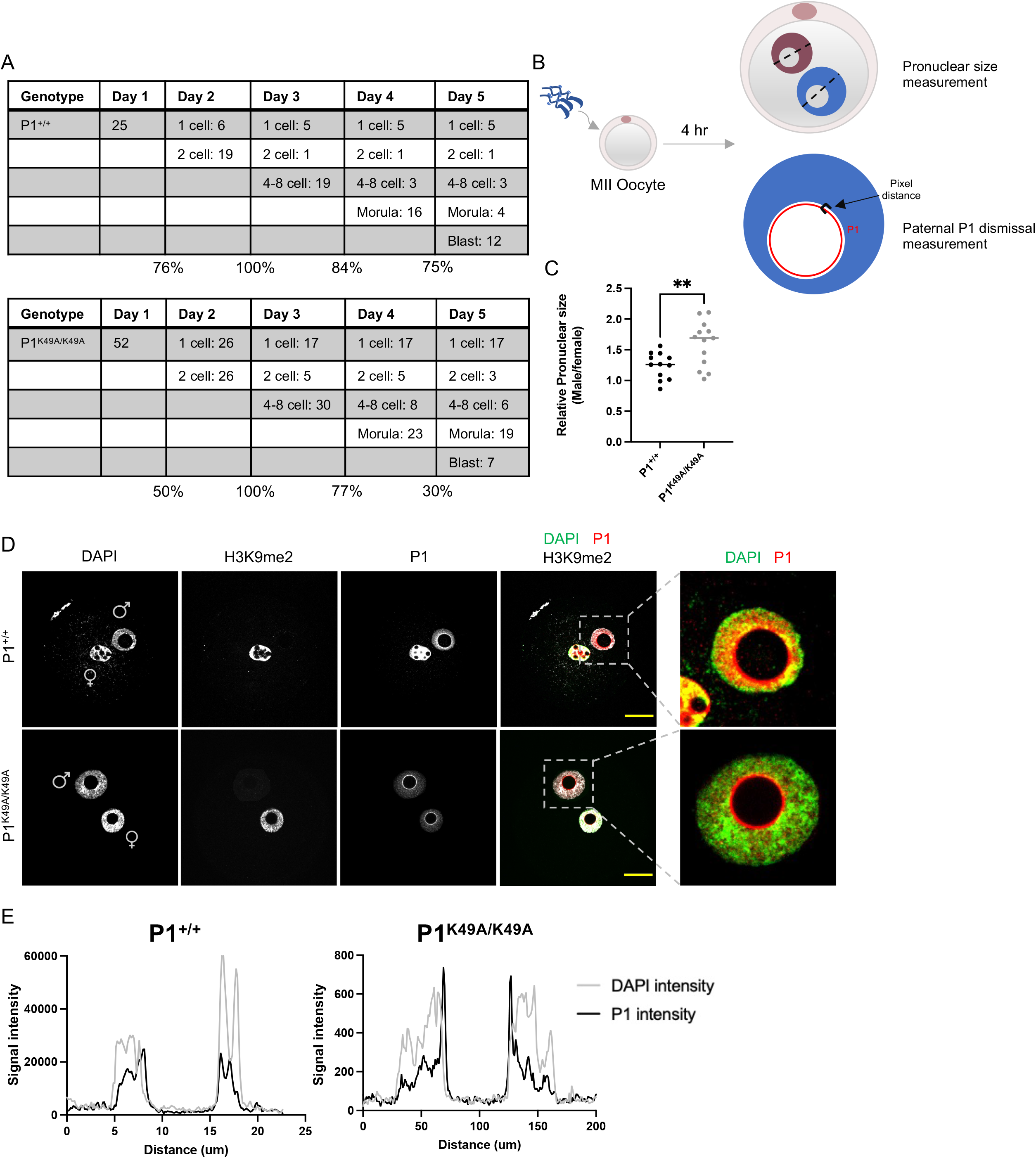
P1 K49A substitution results in decreased blastocyst formation and accelerated P1 dismissal from paternal chromatin. **(A)** Developmental outcomes of intracytoplasmic sperm injections using either P1^+/+^ or P1^K49A/K49A^ sperm reveals a marked decrease in blastocyst formation rate using P1^K49A/K49A^ sperm. Statistical test was performed using a two-sided Fisher’s exact test, p=0.0009. **(B)** Experimental scheme for assessing pronuclear size and P1 removal from paternal chromatin. **(C)** Quantification of relative pronuclear size (male/female) in zygotes derived from either P1^+/+^ or P1^K49A/K49A^ sperm. Statistical test was performed using an unpaired t-test. Quantification was performed from a total of n=12 P1^+/+^ zygotes and n=13 P1^K49A/K49A^ zygotes, p=0.0075. **(D)** Representative immunofluorescence images of zygotes derived from P1^+/+^ or P1^K49A/K49A^ sperm collected at 4hpf and stained for P1. Scale bar = 20 μm.

Since proper decompaction of the paternal genome and the replacement of protamines by histones is critical for embryonic development following fertilization, we first evaluated male pronuclear remodeling and protamine dismissal from paternal chromatin by immunostaining zygotes derived from either P1^+/+^ or P1^K49A/K49A^ sperm (collected 4 hours post-fertilization [hpf], outlined in **Figure 4B**). By DAPI staining alone, we found that the average relative pronuclear size (male/female) in zygotes derived from P1^K49A/K49A^ sperm was significantly larger than in those derived from P1^+/+^ sperm, suggesting potentially an accelerated decompaction of paternal chromatin in the mutant (**Figure 4C**).

If paternal chromatin decompaction is affected by the K49A substitution, we would then expect to observe differences in P1 removal in the zygote. When we stained both P1^+/+^ and P1^K49A/K49A^ zygotes at 4hpf with our custom P1 antibody, we observed differences in P1 distribution patterns. In P1^+/+^ zygotes at 4hpf, P1 is more broadly and densely distributed throughout the male pronucleus and appears more often to be associated with DNA. In contrast, in P1^K49A/K49A^ zygotes, P1 adopts a speckle-like distribution in the male pronucleus and the intensity of a concentrated (possibly phase separated) ring-like pattern immediately inside the prenucleolar body is higher (**Figure 4D**). When we measured the pixel distance from the edge of the DAPI signal to the most intense P1 signal as a proxy of P1 eviction, we found a significantly higher distance for P1^K49A/K49A^ zygotes, further indicating an accelerated P1 dismissal (**Figure 4E, S4A**). Moreover, while ∼24% of P1^+/+^ measurements indicate complete overlap between P1 and DNA (a pixel distance of 0, scale corresponds to ∼6 pixels/μm), only ∼14% of P1^K49A/K49A^ measurements exhibited complete overlap, with 71% of P1^K49A/K49A^ measurements having a pixel distance >3 (compared to 50% of P1^+/+^ zygotes, **Figure S4A**). The increased number of P1^K49A/K49A^ zygotes with premature P1 dismissal is consistent with the higher percentage of arrested 1-cell zygotes in the mutant.

Curiously, while tracking P1 in both WT and mutant zygotes, we noticed that P1 localized to both female and male pronuclei. Moreover, P1 in P1^K49A/K49A^ zygotes displays similar localization patterns in the female pronucleus as the male pronucleus (**Figure 4D, S4C**). Localization of protamines in the female pronucleus was previously reported but was assumed to be an antibody artifact (McLay and Clarke, 2003). To rule out this alternative explanation, we knocked in a V5 tag to the endogenous P1 locus to create an N-terminal V5-tagged P1 protein (V5-P1) and confirmed that the addition of the V5 tag on P1 does not affect sperm parameters or fertility (**Figure S4B**). We detected V5-P1 in both pronuclei using an anti-V5 antibody, consistent with our anti-P1 antibody staining and supporting that localization to the female pronucleus is not an antibody artifact (**Figure S4C**). The protamines detected in the female pronuclei are male derived since a reciprocal IVF (using P1^+/+^ sperm and V5-P1 oocytes) revealed no V5 expression but P1 localization in both pronuclei (data not shown). Although localization to the female pronucleus is an intriguing observation, future studies are required to better understand its potential functional implications.

Altogether, our observations suggest that a single amino acid substitution in P1 results in an increased number of embryos arresting at the 1-cell stage and accelerated P1 removal in P1^K49A/K49A^ zygotes. Thus, these results confirm the functional significance of the K49 residue *in vivo* and at the organismal level.

### The substitution of P1 K49 to alanine decreases P1 DNA binding ability

Given our *in vivo* observations of the P1 K49A substitution and the difficultly of assessing how protamine-DNA binding or dynamics are regulated *in vivo*, we turned to bulk biochemical and single molecule assays *in vitro* to examine protamine-DNA interactions. Because of the high arginine content in protamines, generation of recombinant P1 and P2 proteins in bacteria in sufficient quantity and purity has been extremely challenging. To overcome these challenges, we developed a method to successfully purify P1 and P2 proteins (amino acid sequences shown in **Figure 5A**) from both P1^+/+^ (referred to as WT P1 or WT P2) and P1^K49A/K49A^ mature sperm (referred to as P1 K49A or pro P2) using a combination of acid extraction of basic proteins and size exclusion chromatography (**Figure 5B**). The combination of these two methods enabled purification and efficient separation of P1 and P2 not only from each other but also from other basic proteins such as histones (**Figure S5A**,**B**).

**Figure 5:**
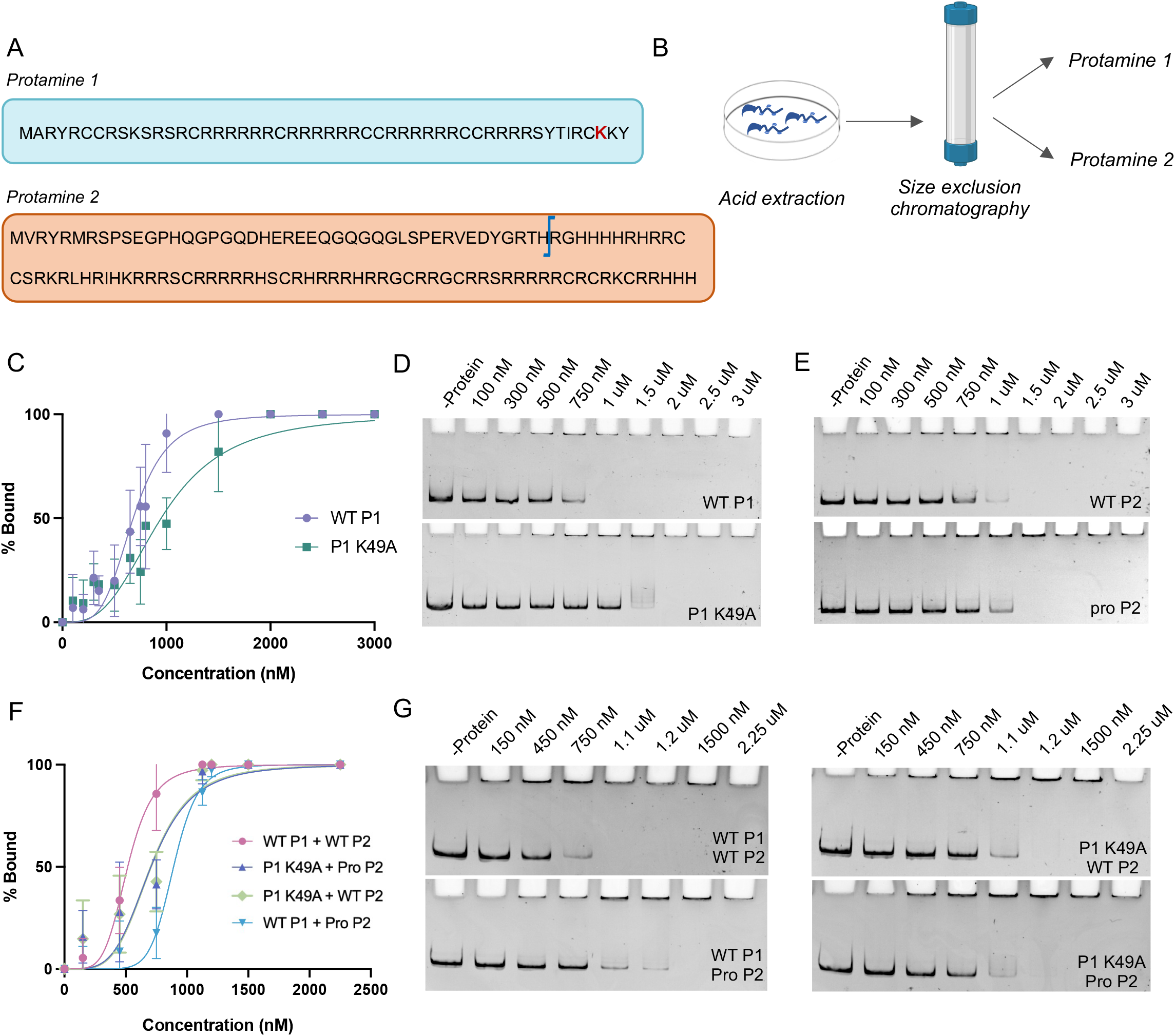
P1 K49A substitution negatively affects DNA binding. **(A)** Schematic of mouse P1 and P2 sequences. Blue bar in P2 indicates cleavage site. **(B)** Purification scheme used for purifying P1 and P2 from mature mouse sperm. **(C)** Quantification of the binding affinities of WT P1 and P1 K49A to a linear ∼300 bp DNA fragment. K_d,app_ values were calculated using the Hill equation and were taken from at least 3 technical replicates per protein. **(D)** Representative EMSAs of a titration of increasing amounts of WT P1 (top) or P1 K49A (bottom) illustrating their interaction with a ∼300 bp DNA fragment. **(E)** Representative EMSAs of a titration of increasing amounts of WT P2 (top) or pro P2 (bottom) illustrating their interaction with a ∼300 bp DNA fragment. **(F)** Quantification of the binding affinities of P1 (either WT or K49A) and P2 (either WT or pro P2) mixed at a 1:2 ratio to a linear ∼300 bp DNA fragment. **(G)** Representative EMSAs of titrations of increasing amounts of indicated P1 and P2 mixed at a 1:2 ratio.

To test the binding affinity of WT P1 or P1 K49A to DNA, we performed electrophoretic mobility shift assays (EMSAs). Using a ∼300 bp linear DNA we found that WT P1 robustly bound DNA in a concentration-dependent manner after 1 hour with a K_d,app_ of 0.68 μM. In contrast, P1 K49A exhibited a marked decrease in DNA binding affinity, K_d,app_ = 0.95 μM (**Figure 5C,D**). Since we noted that the protamine-DNA complex never entered the gel, regardless of experimental conditions, type of gel, or DNA fragment size (data not shown), we inferred that protamine-DNA complexes are forming large, higher order structures that preclude migration into the gel. To confirm this, we repeated the EMSA experiments, in the presence or absence of proteinase K. As expected, the addition of proteinase K dissolved the complex and restored movement of the DNA into the gel, suggesting that the well shift is representative of protamine-DNA interactions and not a technical artifact (**Figure S5H**).

Next, to ensure that the reaction had reached binding equilibrium, we performed EMSAs after incubating DNA and proteins for 10 minutes, 1 hour, or 4 hours, but as we did not observe any difference in binding for either WT P1 or P1 K49A, we concluded that the reaction reaches equilibrium within 10 minutes (**Figure S5D**,**E**). Furthermore, both P1 and P1 K49A appeared capable of cooperative binding behavior, as evidenced by Hill coefficients >1 (4.2 for WT P1 and 3.0 for P1 K49A). The lower values in Hill coefficients in the P1 K49A mutant might reflect a lower DNA affinity or defect in the ability of the mutant to initiate binding and/or polymerize on DNA. To explore whether this apparent cooperative behavior is a general property of protamine proteins or restricted to P1, we also assessed the binding of WT P2 and pro P2. Similarly, WT P2 and pro P2 also displayed cooperative-like behavior *in vitro* (Hill coefficients of 4.2 and 3.7, respectively). Moreover, WT P2 had a higher binding affinity, K_d,app_ = 0.67 μM (similar to that of WT P1), than pro P2, K_d,app_ = 0.84 μM (**Figure 5E, S5C**) and similarly neither binding affinity was affected by incubation time (**Figure S5F**,**G**).

Since mouse sperm (and most mammalian sperm) use both P1 and P2 to package chromatin, we next aimed to understand how the presence of both proteins influences their affinity to DNA, and specifically whether the P1 K49A substitution may alter this affinity. To this end, we repeated the EMSAs using WT P1 or P1 K49A in combination with either WT P2 or pro P2 in a 1:2 ratio (P1:P2), the expected ratio in mouse. As expected, the combination of WT P1 and WT P2 bound more efficiently to DNA than either protein alone (**Figure 5F, 5G top left panel**). However, upon mixing WT P1 with pro P2, we observed a significant decrease in affinity and an overall shift in the binding curve (**Figure 5F, 5G bottom left panel**). When comparing the binding properties of P1 K49A with either WT P2 or pro P2, the binding appeared nearly identical, in both cases requiring a higher protein concentration (∼1.2 uM of total protamine) to reach a fully bound state (**Figure 5F, 5G right panels**).

Taken together, we show that although P1 K49A maintains a cooperative binding mode, the mutant protein has a marked decrease in DNA binding affinity. Furthermore, we show that protamine-DNA binding affinity and cooperative behavior is enhanced in the presence of both P1 and P2 together, but the mutant P1 protein loses its preferred selectivity for mature P2 and instead can interact equally well with either P2 or pro P2.

### P1 K49A substitution causes altered DNA compaction and decompaction kinetics

Since EMSAs ultimately measure an equilibrium between both the protein on rate (k_on_) and off rate (k_off_) and therefore cannot provide kinetic information, we turned to a DNA curtain assay. Here, we used DNA from bacteriophage λ (λ-DNA) to investigate the real time compaction and decompaction kinetics of wild type and mutant protamines at single molecule resolution (**Figure 6A**). Protamines are expected to bind to 10-15 bp sites (Balhorn, 2007) and notably the 50 kb of λ-DNA contains a large diversity of 10-15 bp sites which are all represented thousands of times within the mouse genome, therefore this DNA source allows us to probe representative and relevant general interactions between protamines and DNA. As protamines cause the dissociation of DNA-intercalating dyes such as YOYO-1 (data not shown), we instead labeled each λ-DNA molecule with a fluorescent dCas9 at the untethered end and monitored changes in DNA length to calculate compaction and decompaction (**Figure 6B**).

**Figure 6:**
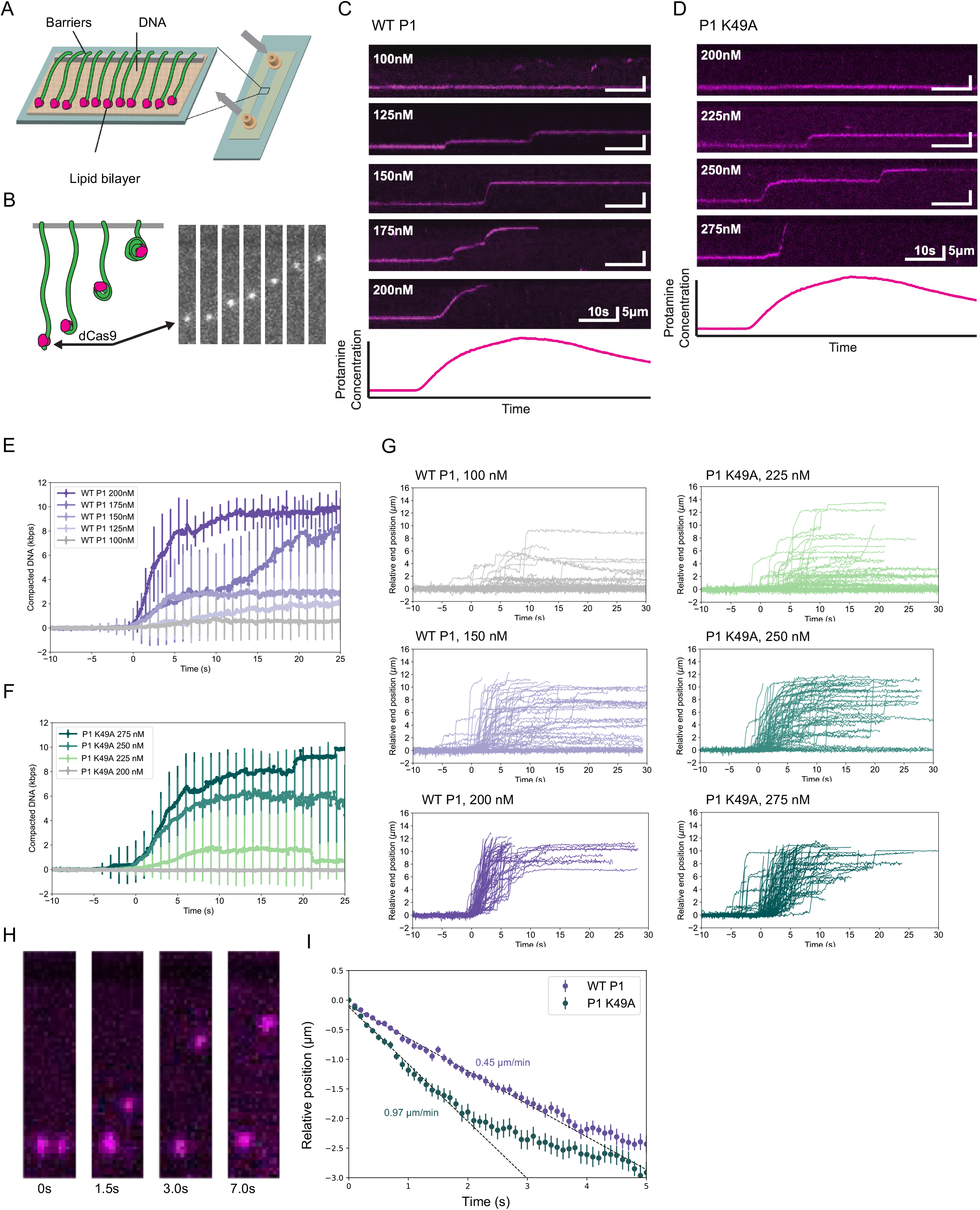
P1 K49A substitution alters DNA compaction and decompaction kinetics *in vitro*. **(A)** Schematic of DNA curtains. DNA molecules are labeled at the 3’ end by dCas9 (shown in pink). **(B)** Cartoon representation shown side-by-side with actual images of protamine-driven DNA compaction. **(C)** Representative kymographs of WT P1 induced DNA compaction at increasing protein concentrations. **(D)** Representative kymographs of P1 K49A induced DNA compaction at increasing protein concentrations. **(E)** Average DNA compaction by WT P1 at increasing concentrations. Error bars represent standard deviations (n=78 traces for 100 nM, n=67 for 125 nM, n=78 for 150 nM, n=63 for 175 nM, and n=66 for 200 nM). **(F)** Average DNA compaction by P1 K49A at increasing concentrations. Error bars represent standard deviations (n=48 traces for 200 nM, n=74 for 225 nM, n=68 for 250 nM, and n=81 for 275 nM). **(G)** Traces of individually tracked DNA molecules over time at low, intermediate, or high concentration of either WT P1 (left panels) or P1 K49A (right panels) illustrating cooperative behavior. **(H)** Representative image of WT P1-driven compaction of adjacent DNA molecules within the curtain highlighting differences in compaction even between DNA molecules that are side-by-side. **(I)** Decompaction of DNA initially compacted by WT P1 and P1 K49A over time illustrates differences in decompaction rates. Error bars represent SEM.

We found that WT P1 largely failed to initiate compaction at 100 nM, but induced robust and complete DNA compaction at 200 nM, with an average velocity of 1.57 μm/s (∼6kbp/s) (**Figure 6C,E,G**). In contrast, P1 K49A failed to initiate compaction at 200 nM but achieved robust compaction at 275 nM (**Figure 6D,G,F**). In addition, the average velocity for P1 K49A at the maximum rate of compaction was slower than WT P1 (1.09 μm/s at 275 nM compared to P1, 1.57 μm/s at 200 nM). Curiously, at the low/intermediate protein concentrations, both WT P1 and P1 K49A displayed a unique pattern: a few molecules condensed >10 kilobases of DNA (**Figure 6G**), but the majority of DNA molecules remained uncompacted, with a few strands only initiating compaction stochastically. At intermediate protein concentrations (125 nM – 175 nM) the extent of DNA compaction is non-uniform (**Figure 6G**), meaning that not all DNA strands within a single experiment compact to the same level and compaction exhibits a start-and-stop behavior (**Figure 6G**). Given that even directly adjacent DNA molecules separated by only a couple microns do not exhibit the same behavior (i.e. one molecule could compact completely, whereas the neighboring DNA molecule does not compact at all, **Figure 6H**), this highlights the need for a better understanding of mechanisms underlying cooperativity and competition in the context of protamine binding, and the local signals that license regions of DNA to compact. Therefore, consistent with the initial cooperative-like behavior we observed in the EMSA experiments, the DNA curtain experiments support that protamines, when present in a limited pool, preferentially bind to a small number of DNA molecules to generate a high level of compaction, as opposed to distributing evenly among all available DNA molecules to produce a uniform but low level of compaction.

Next, we assessed the kinetics of WT P2 and pro P2. Pro P2 initially binds to DNA in elongating spermatids and then undergoes proteolytic cleavage to generate mature P2 (WT P2). To this end, we found that WT P2 required 275 nM protein to achieve robust compaction, similar to P1 K49A, but compacted DNA at twice the rate of P1 K49A at this concentration (2.01 μm/s at 275 nM, **Figure S6A**,**C**). Pro P2 compacts DNA across a similar concentration range but at slightly slower rates (pro P2: 1.26 μm/s vs. WT P2: 1.97 μm/s at 250 nM, **Figure S6B**,**D**,**E**). At intermediate concentrations, and similar to WT and mutant P1, both WT P2 and pro P2 compact a fraction of DNA molecules, leaving many uncompacted (**Figure S6E**). However, the average extent of compaction generated by WT P2 is much greater (e.g. ∼9 μm for WT P2 and ∼4 μm for pro P2 after 5 seconds at 275 nM), consistent with increased genome compaction occurring once P2 has undergone processing (**Figure S6C**,**D**). Unlike for the other tested protamines, we found that WT P2 at intermediate concentrations initiated DNA compaction but then rapidly de-compacted (**Figure S6A**,**C**,**E**). This result suggests that the stability of WT P2-DNA complexes is more sensitive to local concentration changes and initial WT P2-DNA complexes are stabilized by further protein binding. We surmise that these opposing characteristics of WT P2; robust DNA compaction and more sensitive concentration-dependent decompaction, may be central to the opposing roles of protamines in sperm versus zygotes. Overall, the compaction behavior we observe across protamines suggests that tight regulation of local protamine concentration provides a general mechanism for controlling chromatin condensation during spermiogenesis *in vivo*.

The DNA curtain experiments also allowed us to monitor DNA decompaction, the rate at which protamines passively dissociate from DNA. Here, we find that DNA condensed by P1 K49A decompacted significantly faster than DNA decompacted by WT P1 (0.97 μm/min, ∼3.6 kbp/min for P1 K49A vs. 0.45 μm/min, ∼1.7 kbp/min for WT P1) (**Figure 6I**). Likewise, pro P2-compacted DNA decompacted at a faster rate than DNA stably compacted by WT P2 (0.45 μm/min, ∼1.7 kbp/min vs. 0.37 μm/min, ∼1.4 kbp/min) (**Figure S6F**). In short, these data demonstrate that mutant P1, even at higher concentrations relative to WT P1, compacts DNA slower and dissociates from DNA faster, consistent with our bulk measurements of K_d,app_. Furthermore, compared to pro P2, WT P2 requires more protein to initiate compaction, but compacts DNA at a faster rate and to a greater extent, and dissociates from DNA more slowly, again consistent with our bulk results. Moreover, our *in vitro* studies also show that P1 K49A-compacted DNA decompacts significantly faster than DNA compacted by WT P1, explaining our observation in embryos. Therefore, although electrostatics may be a prominent driver of sperm DNA condensation, other regulatory factors, like PTMs or individual residues, fine-tune the DNA compaction and decompaction to ensure correct packaging and developmental sequence of events.

## Discussion

Efficient eviction of histones and subsequent addition of protamines to optimally package paternal chromatin during spermiogenesis is essential to safeguard fertility throughout life. Given the arginine-rich composition of protamines, previous work assumed a non-specific protamine-DNA binding mechanism, leaving the contribution of individual P1 or P2 residues to chromatin condensation in spermatids unresolved. Here, we pioneered a series of complementary molecular, genetic, biochemical, and biophysical assays to explore how the single amino acid substitution of P1 K49 to an alanine—a residue outside the central arginine core—systematically perturbs sperm genome packaging. Our systematic *in vitro* and *in vivo* analysis of efficacy, development, and biophysical properties of the P1 protein support a possible regulatory role for the K49 residue. Based on our findings, we propose a reevaluation of the conventional view of protamines as purely electrostatic structural components to instead consider that protamine protein sequence variants outside of the arginine core may have evolved to execute species-specific and regulated packaging and unpackaging processes.

The conservation of K49 in P1 across the rodent lineage led us to hypothesize that it plays an essential and species-specific role in spermiogenesis and/or embryonic development, either through the K49 residue itself or through its acetylation. By using a modification-specific antibody against K49ac, we find that acetylation is acquired in early elongating spermatids, and that the K49A substitution leads to a ∼3.5-fold increase in canonical histone retention and accumulation of pro P2. Given that the K49A substitution affects P1-DNA binding affinity, more P1 K49A protein is likely needed *in vivo* to overcome this decrease in affinity. Hence, it is possible that the efficiency of histone eviction is secondarily hindered, resulting in increased histone retention. Earlier studies have also elegantly shown that histone acetylation is a prerequisite for spermatid maturation and histone-to-protamine exchange, and furthermore, disruption of the testes-specific dual bromodomain containing protein, BRDT—an acetyl-lysine reader—also precludes nucleosome eviction in round spermatids (Boussouar et al., 2014; Dong et al., 2017; Gaucher et al., 2012; Luense et al., 2019). Here, we show that the P1 K49A mutation alters the site of modification – preventing acetylation on P1. It is possible that P1 acetylation is somehow read by BRDT and is involved with nucleosome eviction or retention. Our findings suggest that P1 K49ac may be important in the histone-to-protamine exchange process, thereby expanding the pool of factors implicated in this process. Furthermore, given that both bromodomains of BRDT are required to induce a large-scale acetylation-dependent chromatin reorganization in sperm, our future studies will explore the possibility that BRDT may interact with both acetylated histone H4 and acetylated P1 to modulate this process.

In addition to altered sperm chromatin composition, we find that fertility in P1^K49A/K49A^ males is significantly decreased due to a near-total loss of progressive sperm motility. The morphological defects we observed in P1^K49A/K49A^ sperm overlap largely with those observed in mice that are haploinsufficient for P1 (P1^+/-^), lack P2, or lack TNP1 and TNP2 (Schneider et al., 2016; Shirley et al., 2004b; Takeda et al., 2016b; Yu et al., 2000b). Interestingly, many of the morphological abnormalities localize to the midpiece (sperm head/neck connection), raising the possibility that protamine incorporation is linked to cytoskeleton remodeling or manchette formation in elongating spermatids. This conclusion is also supported by the previously reported interaction between phosphorylated P1 and the inner nuclear membrane protein Lamin B receptor (Mylonis et al., 2004). Hence, active cytoskeletal remodeling in the setting of nuclear remodeling in sperm could be a cellular mechanism that overrides the repulsive forces of positively charged proteins, such as protamines. Conversely, this remodeling process may increase local protamine concentration and enhance protamine-DNA cooperativity to ensure regulated initiation and polymerization of protamines on genomic segments—potentially ensuring that sperm DNA compaction is able to reach an energetically favorable structure while also maintaining instructive information for programmatic unfolding during development.

The large net positive charge of arginine-rich protamines remains a challenge to a more complex model of protamine behavior and regulation, as their interaction with DNA is undoubtedly electrostatic. Our data suggest that electrostatic interactions are important but are not the only determinants of protamine-DNA interactions. The differences in P1 DNA binding affinity we observed in protamines across species suggest a possible role for protamine sequence, protein-protein oligomerization, or possibly differences in protamine PTMs (subjects of future investigations). For example, *in vitro*, we observe that P1 K49A has a much lower binding affinity for DNA than WT P1. This impact from the loss of a single lysine residue is unexpected, bearing in mind that this protein contains more than 30 positively charged residues (**Figure 5C,D, 6C-F**) (Berg and von Hippel, 1987; Shultzaberger et al., 2007). Similarly, in the DNA curtain assay, we found that each protein displayed a sharp concentration dependence on the level of DNA compaction measured on DNA curtains. Moreover, compaction was not uniform across the curtain as was observed for DNA compaction by HP1 (Larson et al., 2017). Remarkably, even for DNA molecules side-by-side within our experiments, we observed drastically different levels of compaction when incubated with protamines, indicative of distinct levels of protamine binding on a molecule-to-molecule basis. Furthermore, small differences in protein concentration resulted in unexpectedly large changes in compaction velocity. For example, increasing WT P1 from 175 nM to 200 nM resulted in an increase in average velocity from 0.31 μm/s to 1.57 μm/s. Taken together, we hypothesize that protamines engage in cooperative binding modes with DNA and that tight regulation of their local concentration is a mechanism for achieving precise control over chromatin condensation during spermiogenesis *in vivo*. In addition to differences in compaction rates, the P1 K49A protein dissociated from DNA significantly faster than DNA compacted by WT P1. Strikingly, this result is in agreement with the accelerated dismissal of P1 from the paternal genome in zygotes (**Figure 4D, S4D**) and suggests that modifications apart from phosphorylation may help to regulate the protamine-to-histone transition following fertilization.

Altogether, our findings highlight an indispensable role for P1 K49 in protamine biology, as illustrated by the significant perturbations in protamine-DNA interactions, sperm chromatin packaging, and embryonic development that occur when substituting K49 for alanine. Furthermore, these results highlight the potential that other amino acid residues in P1 outside the central arginine-rich DNA binding core may have functional consequences and perturb biological processes. Our observation that a single amino acid substitution can cause such dramatic alterations in the biophysical and functional properties of mouse P1 lends strong support to the conclusion that evolutionary changes in protamine protein sequences across species are unlikely to be neutral. Future studies are needed to test whether additional residues or PTMs have an impact on fertility. Curiously, sites of modification in both mouse and human protamines are enriched in the N- and C-terminal sequences flanking the arginine core and tend to be highly conserved within a species, but not across species. Therefore, it is conceivable that such non-arginine residues evolved to regulate the species-specific sperm genome packaging and subsequent unpackaging in the zygote to ensure both species compatibility upon fertilization and optimal organismal reproductive fitness.

## Author Contributions

S.S.H., L.M., and S.B.S. provided overall project design. L.M., S.B.S., M.R., and C.S. performed experiments. Y.S. performed ICSI experiments. D.P. and R.A. assisted with purification of protamines using chromatography for *in vitro* biochemistry. K. J. constructed/analyzed phylogenetic trees and performed embryo quantifications. S.R. performed DNA curtain experiments. L.M. and S.S.H. wrote the manuscript with input from S.R. Comments from all authors were provided.

## Acknowledgements

We thank members of the Hammoud Lab for scientific discussions and manuscript comments; Dr. Yueh-Chiang Hu and members of the Cincinnati Children’s Hospital Transgenic Animal and Genome Editing Core Facility; Dr. Thomas Saunders and members of the University of Michigan Transgenic Animal Model Core. Portions of Figure 7 were created with BioRender.com. This research was supported by National Institute of Health (NIH) grants 1R21HD090371-01A1 (S.S.H), 1DP2HD091949-01 (S.S.H.), R01 HD104680 01 (S.S.H), 5K12 HD065257-07 (S.B.S.), 1R03HD10150101A1 (S.B.S), R01-AG050509 (J.N.), R01-GM120094 (J.N.), training grants NSF 1256260 DGE (L.M.), Rackham Predoctoral Fellowship (L.M. and X.Z.), T32GM007315 (L.M.), an American Cancer Society Research Scholar grant RSG-17-037-01-DMC (J.N.), an American Heart Association predoctoral fellowship award ID: 830111 (R.A.), and Open Philanthropy Grant 2019-199327 (5384) (S.S.H.).

---

**Figure 7:**
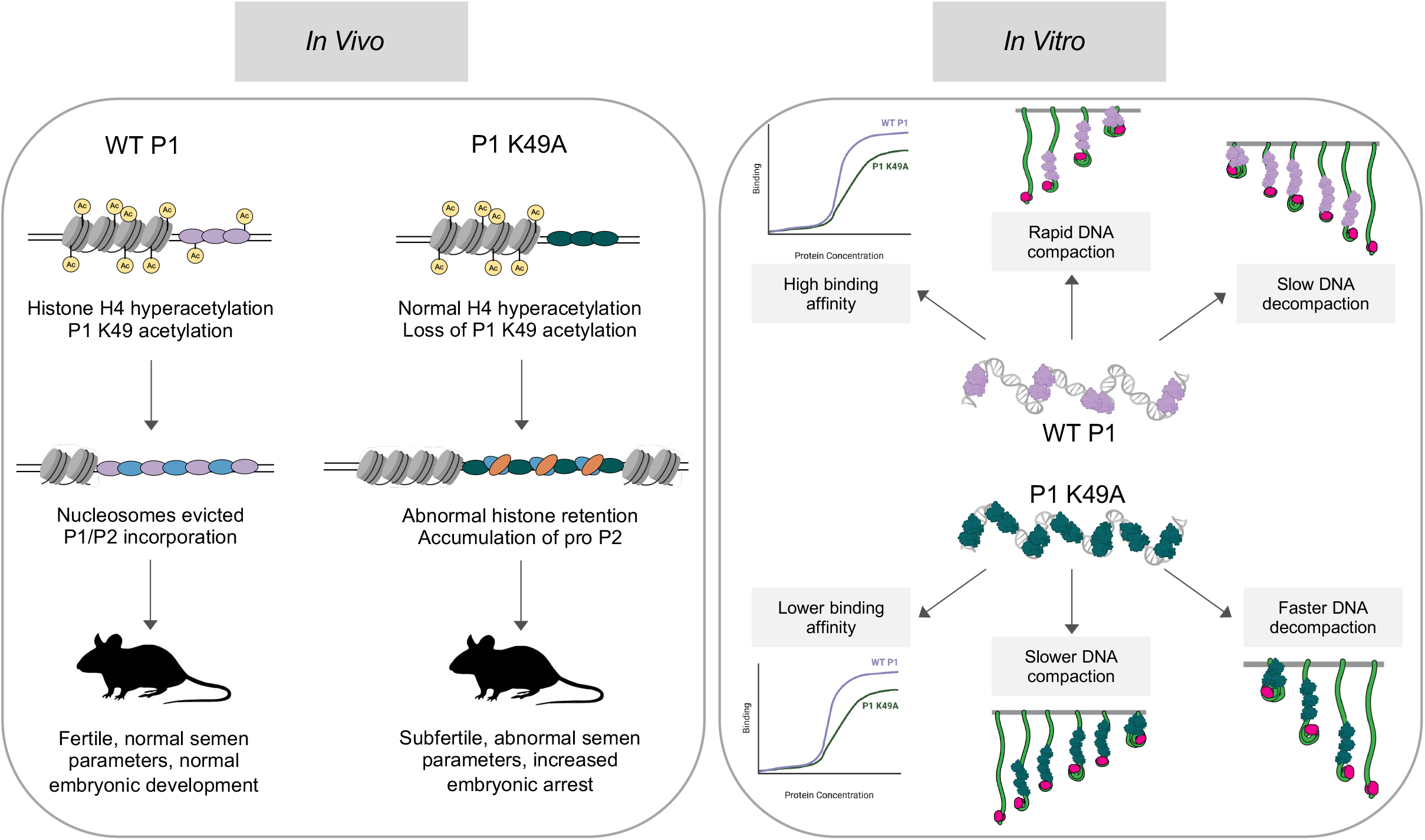
Model of alterations to both *in vivo* and *in vitro* protamine functions caused by P1 K49A substitution.

## Methods

### RESOURCE AVAILABILITY

#### Lead Contact

Additional information and requests for resources and reagents should be directed to and will be fulfilled by the Lead Contact, Saher Sue Hammoud.

#### Materials Availability

All unique reagents generated in this study are available from the Lead Contact with a completed Materials Transfer Agreement.

#### Data and code availability

All data reported in this paper can be shared by the lead contact upon request. No unique codes were generated in this study.

### EXPERIMENTAL MODEL AND SUBJECT DETAILS

#### Mice

All experiments using animals were carried out with prior approval of the University of Michigan Institutional Committee on Use and Care of Animals (Protocols: PRO00006047, PRO00008135, PRO00010000) and in accordance with the guidelines established by the National Research Council Guide for the Care and Use of Laboratory Animals. Mice were housed in the University of Michigan animal facility, in an environment controlled for light (12 hours on/off) and temperature (21 to 23°C) with ad libitum access to water and food (Lab Diet #5008 for breeding mice, #5LOD for non-breeding animals).

P1^K49A/K49A^ knock-in mice were generated on the C57BL/6N background using CRISPR/Cas9-mediated genome editing by the Cincinnati Children’s Hospital Transgenic Animal and Genome Editing Core Facility. The sgRNA and donor oligo were designed as previously described (Haeussler et al., 2016; Yuan and Hu, 2017). The guide RNA target sequence was selected according to the on- and off-target scores provided by the web tool CRISPOR (Haeussler et al., 2016) (http://crispor.tefor.net) and proximity to the target site. Ribonucleoprotein (RNP) complexes were formed by mixing the sgRNA (80 ng/uL) with Cas9 protein (IDT, 120 ng/uL) in Opti-MEM (ThermoFisher) and incubating at 37 °C for 10 minutes, at which time the donor oligo (IDT, 500 ng/uL) containing the intended mutation was added. Zygotes from super-ovulated C57BL/6N females were electroporated with 7 uL of the RNP/donor oligo mix on ice using a Genome Editor electroporator (BEX; 30V, 1 ms width, 5 pulses with 1 s interval). Two minutes after electroporation, zygotes were moved to 500 ul cold M2 medium (Sigma), warmed to room temperature, and transferred to oviductal ampullas of pseudopregnant CD-1 females. All animal procedures were carried out in accordance with the Institutional Animal Care and Use Committee and approved protocol of Cincinnati Children’s Hospital Medical Center. Offspring were genotyped for the P1 K49A mutation by extraction of genomic DNA from a small ear biopsy. Mutant males and control mice were used for all experiments between 8-16 weeks of age for all studies.

### METHOD DETAILS

#### Antibodies

Rabbit polyclonal antibodies against total P1 and acetylation at P1 K49 were generated at GeneMed Synthesis Inc. via immunization of rabbits with the following synthesized peptides: P1-CRRRRSYTIRSKKY, P1 K49ac-CRRRRSYTIRCK(ac)KY. All other antibodies used are provided in the Key Resources Table.

#### Acid extraction of sperm basic proteins

Extraction of basic proteins from mature sperm was performed as previously described (de Yebra and Oliva, 1993). Briefly, sperm pellets were subjected to hypotonic lysis in 1 mM PMSF and subsequently spun down at 8,000xg for 8 min. Sperm pellets were then resuspended in 100 uL of 100 mM Tris pH 8.0, 20 mM EDTA, and 1 mM PMSF followed by denaturation of proteins with 100 uL of 6 M Guanidine-HCl, 575 mM DTT and alkylation with 200 uL of 522 mM sodium iodoacetate for 30 min in the dark. Protein pellets were then washed twice with 1 ml ice cold ethanol and extracted with 800 uL of 0.5 M HCl, 50 mM DTT at 37°C for 10 min. Supernatants were precipitated overnight at -20°C with TCA to a final concentration of 20%. The following day, precipitates were spun down at 12,000xg for 8 minutes and protein pellets were washed twice in 1 ml of 1% 2-mercaptoethanol in cold acetone. Final protein pellets were then resuspended in water.

#### Peptide competition assay to assess antibody specificity

Protamines were first acid-extracted using the method described above. An increasing amount of protein (0.5 ug and 1 ug for non-specific peptide, 1 ug and 3 ug for non-acetylated P1 peptide) was loaded on each immunoblot. Prior to adding to immunoblots, antibodies were incubated at room temperature for 30 minutes, with either 10-fold excess of specific or non-specific peptide, or alone with no peptide. After blocking, blots were then incubated for 1.5 hours in either antibody only, antibody with specific peptide, or antibody with non-specific peptide. The non-specific peptide used in these assays was N-DSNKEFGTSNESTE-C and the non-acetylated P1 peptide used was N-CRRRRSYTIRSKKY-C.

#### Mass spectrometry analysis of mouse protamines

Mass spectrometry of mouse sperm was performed at MS BioWorks in Ann Arbor, MI. Briefly, Protamines were first acid-extracted using the method described above. Approximately 20 ug of acid-extracted protein was run in triplicate on a 4-20% SDS-PAGE gel (BioRad) and a single band corresponding to P1 and P2 was cut out for processing. Gel bands were washed once with 25 mM ammonium bicarbonate followed by three washes in 100% acetonitrile. Bands were then reduced with 10 mM dithiothreitol at 60 °C followed by alkylation with 50 mM light iodoacetamide at room temperature. Bands were then digested with either trypsin (Promega) at 37 °C for 4 hours, Chymotrypsin (Promega) at 37 °C for 12 hours, or Lys-C (Promega) at 37 °C for 12 hours. For all enzymes used, digests were quenched with formic acid and the supernatants were analyzed. Digests were analyzed by nano LC/MS/MS with a Waters NanoAcquity HPLC system interfaced to a ThermoFisher Q Exactive. Peptides were loaded on a trapping column and eluted over a 75 um analytical column at 350 nL/min. Both columns were packed with Luna C18 resin (Phenomenex). The mass spectrometer was operated in data-dependent mode, with MS and MS/MS performed in the Orbitrap at 70,000 FWHM and 17,500 FWHM resolution, respectively. The fifteen most abundant ions were selected for MS/MS. Data were searched using a local copy of Byonic with the following parameters: Enzyme: Semi-Trypsin or None (for Chymotrypsin and Lys-C), Database: Swissprot Mouse (forward and reverse appended with common), fixed modification: carbamidomethyl (C), variable modifications: oxidation (O), acetyl (protein N-term), deamidation (NQ), phosphor (STY), methyl (KR), dimethyl (KR), trimethyl (K), mass values: monoisotopic, peptide mass tolerance (10 ppm), fragment mass tolerance (0.02 Da), max missed cleavages: 2. Mascot DAT files were parsed into the Scaffold software for validation, filtering and to create a non-redundant list per sample. Data were filtered using a minimum protein value of 95%, a minimum peptide value of 50% (Prophet scores) and requiring at least two unique peptides per protein. Site localization probabilities were assigned using A-Score (Beausoleil et al., 2006).

#### Evolutionary analysis of P1 sequence conservation

The phylogenetic tree was constructed using maximum likelihood (PhyML) inferred from P1 protein sequences for species across the orders Rodentia, Primate, and Artiodactyla, using the Whelan and Goldman matrix (WAG) substitution strategy. Sequences were downloaded from NCBI and aligned using standard parameters of MUSCLE (Edgar, 2004). Bootstrap support with 1,000 replicates is shown for each node, with values >95 indicating strong support.

#### Immunofluorescence and quantification of seminiferous tubule staging

Adult testes were fixed overnight in 4% PFA at 4°C before submerging in ethanol and processing for formalin fixed paraffin embedding (FFPE). Five-micron thick tissue sections were first deparaffinized followed by permeabilization and subsequent antigen retrieval via boiling in 10 mM sodium citrate pH 6.0 for 10 minutes. Following blocking in 1X PBS, 3% BSA, 500 mM glycine, sections were incubated with primary antibodies overnight at 4°C. PNA-Lectin (GeneTex) was used to stain acrosomes and DAPI was used as a nuclear counterstain. All AlexaFlour-conjugated secondary antibodies (Life Technologies/Molecular Probes) were used at 1:1000. For assessment of staging, seminiferous tubules were split into categories (I-III, IV-VI, VII-VIII, IX, X-XI, XII) according to their Lectin staining pattern and cell types present as previously described (Nakata et al., 2015).

#### Phenotypic assessment of P1^+/+^, P1^K49A/+^, and P1^K49A/K49A^ males

All phenotyping was carried out in males between 64 and 71 days of age (9-10 weeks). Sperm were counted using a Makler chamber and performed as n=3 independent technical replicates per mouse (n=4 mice per genotype). For progressive sperm motility assessment, a minimum of 100 total sperm were counted and forward (progressive) movement was assessed in comparison to the total number of sperm counted in a total of n=4 mice per genotype. For quantification of fecundity, 8-week-old males (n=3 per genotype) were individually housed for 3 days before 8-week-old C57BL/6J females were added. Females (n=3 females per male, for a total of n=9 females per genotype of male) were checked daily for the presence of copulatory plugs and once plugs were noted, females were removed and placed in a new cage. The percent of females that were successfully impregnated was recorded (fecundity).

#### Acid urea gel electrophoresis for the separation of sperm basic proteins

Protamines were first acid-extracted from a fixed number of sperm cells per genotype as described above, with slight modification (Giorgini et al., 2002). Following hypotonic lysis in 1 mM PMSF, sperm pellets were resuspended in 1 ml of 6 M Guanidine-HCl, 500 mM Hepes pH 7.5, 10 mM DTT for 1 hour at room temperature. Cysteines were then alkylated using vinylpyridine to a final concentration of 250 mM and incubated for 1.5 hours at room temperature. Proteins were then extracted with 0.9 M HCl and dialyzed overnight at 4°C against 0.2 M HCl. The following day, insoluble proteins were removed by centrifugation at 12,000xg for 5 minutes. Soluble proteins were then precipitated with TCA to a final concentration of 20% for 4 hours at -20°C. Precipitated proteins were then washed twice with acetone before being resuspended in 0.9 M acetic acid, 8 M urea, 100 mM betamercaptoethanol. Acid urea gels were prepared as previously described (de Yebra and Oliva, 1993). As an identical number of input sperm was used for extraction, an identical volume of protein was loaded for each genotype. P1:P2 ratios were calculated using ImageJ.

#### Sperm protein extraction for the assessment of histone retention

Histone levels in P1^+/+^, P1^K49A/+^, and P1^K49A/K49A^ sperm (sperm pooled from 2-3 animals per genotype) were assessed as previously described (Luense et al., 2019). Briefly, sperm pellets were resuspended in lysis buffer (20 mM Tris pH 7.5, 1 mM MgCl_2_, 1 mM CaCl_2_, 137 mM NaCl, 10% glycerol, 1% NP-40, 12.5 U/ml Benzonase, 1X protease inhibitors), sonicated briefly, and rotated for 1 hour at 4°C. For immunoblotting, lanes were loaded by input number of sperm cells. Due to variability between antibodies, the following sperm numbers were loaded for each corresponding antibody: histone H3, histone H2B, ac-H4, H4K20me3, and H3K9me2: 25000, 50000,10000; histone H4: 100000, 250000, 400000; H3K27ac and H3K27me3: 250000, 500000, 750000. Each blot was probed for alpha tubulin as a loading control to ensure comparative loading.

#### Protamine purification and *in vitro* electrophoretic mobility shift assays

Acid-extraction of sperm basic proteins was first performed as described above. Following TCA precipitation, the protein pellet was resuspended in 50 ul of water and brought up to 500 ul in gel filtration buffer (25 mM Hepes pH 7.5, 150 mM NaCl, 5 mM TCEP (TCEP was not pH neutralized). The solution was then subsequently subjected to size exclusion chromatography using a Superdex S75 column. Peak fractions were identified by absorbance at 214 nm and confirmed by immunoblotting. For *in vitro* electrophoretic mobility shift assays, varying concentrations of purified protamines were incubated with 40 nM DNA (280 bp) after briefly incubating the proteins at 37°C for 10 minutes in reaction buffer. DNA was prepared by PCR amplification of mouse genomic DNA using the primers specified in Supplemental Table 3. After 1 hour of incubation, EMSA reactions were then run on a non-denaturing 0.5X TBE 6% polyacrylamide gel and stained with ethidium bromide (Sigma). Band intensities were quantified using ImageJ.

#### DNA Curtains

Microfluidic devices were constructed, and DNA curtain assays were performed as previously described (Gallardo et al., 2015; Larson et al., 2017). Briefly, a lipid bilayer was coated on the surface of the sample chamber and biotinylated phage DNA was anchored to biotinylated lipids within the bilayer via streptavidin. DNA were then aligned at microfabricated barriers using buffer flow. In all experiments, care was taken to ensure that DNA molecules were separated by at least 2μm to prevent protamine interactions across DNA molecules.

#### DNA labeling with Cas9

Recombinant dCas9 protein was purchased from IDT (Alt-R S.p. dCas9 protein V3). The dCas9 protein was loaded with a dual guide RNA according to IDT’s “Alt-R CRISPR-Cas9 system - in vitro cleavage of target DNA with RNP complex” protocol with slight modifications for DNA curtains. dCas9 protein was diluted to 200nM and incubated with gRNA targeting position 47,752 (AUCUGCUGAUGAUCCCUCCG) at a 1:10 ratio in imaging buffer (1mg/mL BSA, 40mM Tris-HCl, pH=7.5, 50 mM NaCl, 5mM MgCl2, and 1mM DTT) and incubated on ice for at least 15min. Then, anti-HisAlexa555 (Invitrogen) was added to the reaction at a 1:2 ratio to dCas9 and incubated in the dark for 15min at room temperature. The fluorescent RNP complex was then diluted to 4nM in imaging buffer and incubated in the flow cell with DNA for 10 minutes. Finally, to remove nonspecifically bound proteins, the flow cell was washed with 500uL of imaging buffer containing 100ug/mL Heparin.

#### DNA compaction and decompaction experiments

DNA were maintained in flow at a rate of 0.6mL/min (average extension to 90% of contour length) for the duration of compaction and decompaction experiments. Prior to introduction into the flowcell, protamines were incubated at 37°C for 15 min in imaging buffer. Then, protamine was injected into the flowcell, images were collected at 10Hz, and compaction was monitored by tracking the motion of dCas9 molecules. Immediately following compaction, collection was shifted to 0.2Hz and decompaction was monitored by tracking the position of dCas9 molecules.

#### Intracytoplasmic sperm injections and embryo immunofluorescence

Oocyte collection, sperm collection, and piezo-actuated intracytoplasmic sperm injections were performed as previously described (Yoshida and Perry, 2007). Embryos were cultured for 4 hours in KSOM (Millipore-Sigma) prior to immunofluorescence analysis.

For immunofluorescence, embryos were collected at the indicated timepoints after washing in KSOM, treated briefly (30 seconds-1 minute) with Acidic Tyrodes solution (EMD Millipore) to remove the zona pellucida, and fixed in 4% PFA for 10 minutes. Embryos were washed and gently permeabilized in PBS supplemented with 0.1% Triton and 3% BSA overnight at 4 °C before a 1-hour permeabilization in PBS supplemented with 0.5% Triton and 3% BSA. Embryos were then blocked in PBS with 0.1% Triton, 3% BSA, and 10% fetal bovine serum for 1 hour and stained with primary antibodies overnight in blocking buffer at 4°C. The following day, embryos were washed in PBS with 0.1% Triton and 3% BSA five times for 15 minutes each, followed by incubation in secondary antibodies (Life Technologies/Molecular Probes) and DAPI (Sigma) for 2 hours at room temperature. All images were taken on a Nikon A1R-HD25 confocal microscope and processed with ImageJ.

### QUANTIFICATION AND STATISTICAL ANALYSIS

All statistical analyses were performed using the Prism software. Statistical details including the exact statistical test used, exact value of n, what n represents, dispersion measures, and significance values are presented in each corresponding figure legend.

## Supplemental Table Legends

**Table S1. Mass spectrometry analysis of mouse protamine PTMs**

A. Peptides identified from mass spectrometry analysis of mouse P1 and corresponding quality metrics, including localization probability and Ascore.

**Table S2. Synthetic peptides used in this study**

A. List of all peptides used, including the name and sequence.

**Table S3. Oligonucleotides used in this study**

A. List of all oligonucleotides used, including the name and sequence.

