## Supplemental figures for "Single residue substitution in protamine 1 disrupts sperm genome packaging and embryonic development in mice"

### Supplemental Figure 1

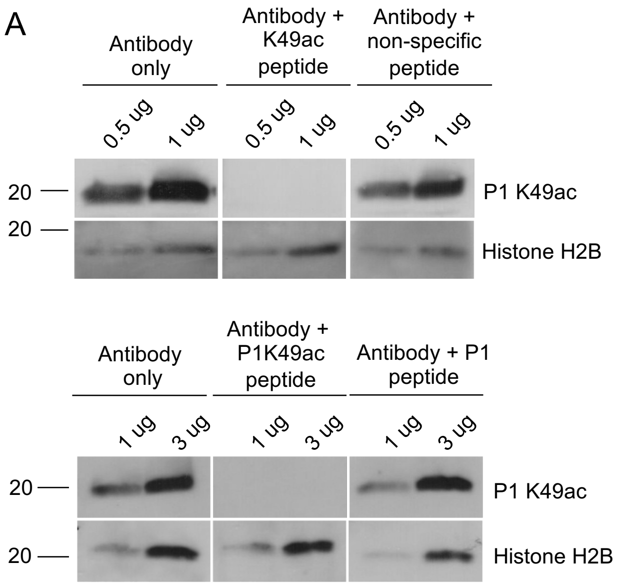


**Supplemental Figure 1: P1 lysine 49 acetylation is acquired in the testis in a stage-specific manner and is present in mouse sperm**. **(A)** Immunoblot of acid extracted protein lysates from mature sperm illustrates a clear band for P1 K49ac that is competed off only in the presence of a specific peptide containing acetylated P1 at K49. Top western blot was performed using a non-specific peptide from an unrelated protein and bottom western blot was performed using a P1 non-acetylated peptide. **(B)** Immunofluorescence of adult testes cross sections using pre-immune serum from rabbits used for antibody generation (top panels) or secondary antibody only (bottom panels) further illustrates specificity of the antibody. Scale bar: 50 μm.

### Supplemental Figure 2

A

| Genomic location | Number of mismatches | Sequence (including mismatches) | Genomic location |
|---|---|---|---|
| Chr6: 129535854 | 0 | GCAGTGGCTCATACACCATAGGG | Intergenic |
| Chr12: 27066405 | 0 | GCTACCACTCTTACACCATAGGG | Intergenic |
| Chr12: 5373801 | 0 | GCCGCCTCGCAAACACCATAGGG | Intron: Klhl29 |
| Chr10: 129362223 | 0 | GCTGTCGATAATACACCATAGAG | Intergenic |
| Chr14: 102981772 | 0 | ACCGCGGCTCCTACACCATCGGG | Exon: Kctd12 |
| Chr6: 72119256 | 0 | GCAGCCTCTCCTACACAATAAGG | Intron: St3gal5 |
| Chr1: 167267189 | 0 | GCCCCCTCCCATACACCACAGGG | Intron: Uck2 |
| Chr2: 168467316 | 0 | GCCACCACTCAGACACCAGATGG | Intergenic |
| Chr12: 4868024 | 0 | GTCGCCCCTCATCCACAATAAGG | Intron: Mfsd2b |

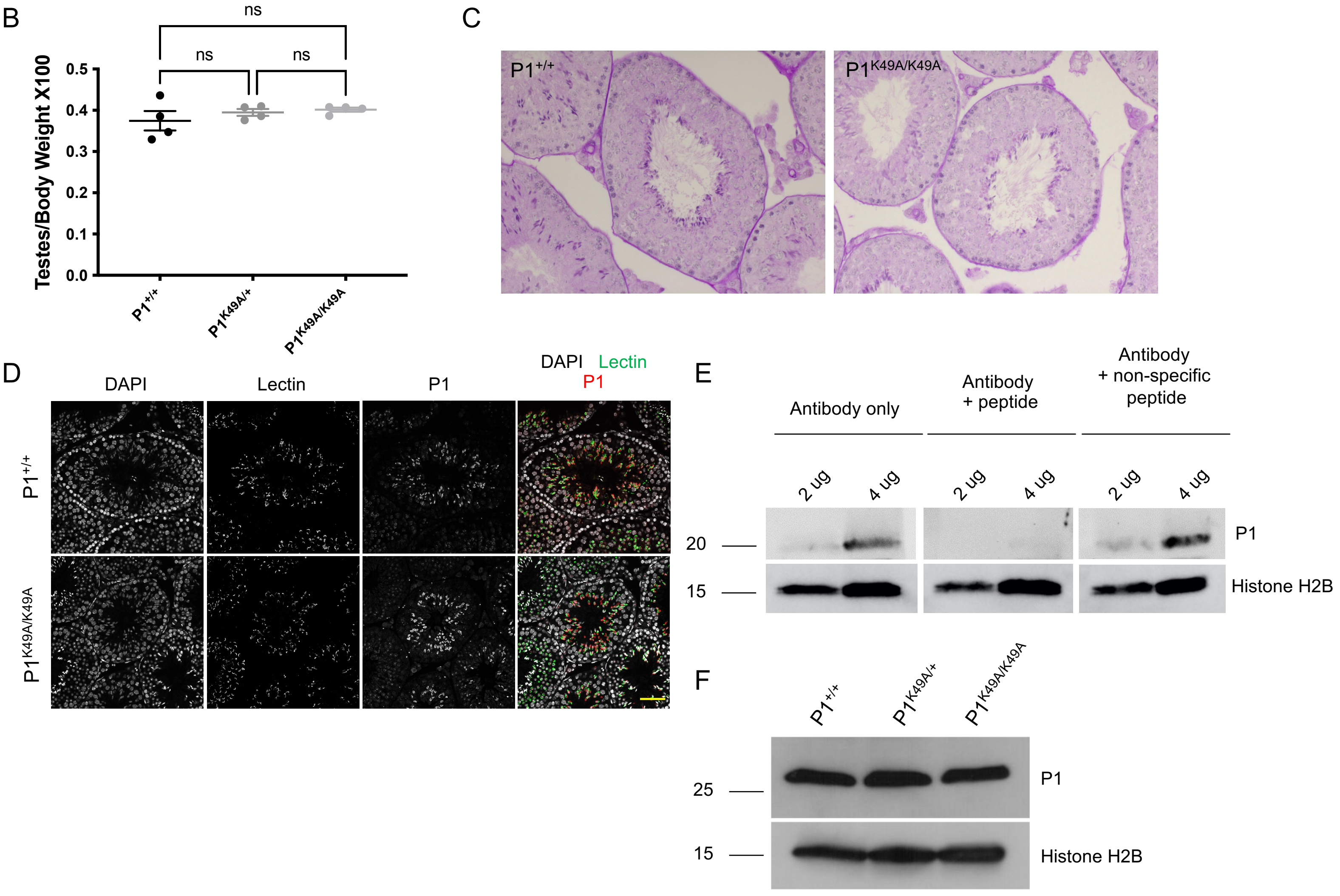


**Supplemental Figure 2: P1 K49A substitution results in sperm motility defects and subfertility.** **(A)** List of potential off-targets and corresponding sequencing results verify no off-target modifications generated by CRISPR/Cas9 editing. **(B)** Testes/body weight ratio of $P1^{+/+}$, $P1^{K49A/+}$, and $P1^{K49A/K49A}$ males (n=4 per genotype) suggests no loss of germ cell populations due to P1 K49A substitution. Statistical test was performed using a one-way ANOVA and adjusted for multiple comparisons, p=0.4391. **(C)** Periodic acid Schiff (PAS)-stained adult testes cross sections highlights normal testis morphology in $P1^{K49A/K49A}$ males. **(D)** Immunofluorescence of adult testes cross sections from $P1^{+/+}$ and $P1^{K49A/K49A}$ testes stained for P1 shows no loss of P1 expression upon substitution of K49 for alanine. Scale bars: 50 µm. **(E)** Peptide competition immunoblot for custom P1 antibody highlights antibody specificity. **(F)** Immunoblot of RIPA-extracted testes from $P1^{+/+}$, $P1^{K49A/+}$, and $P1^{K49A/K49A}$ males shows comparable expression of P1 across all genotypes.

**Supplementary Figure 3**

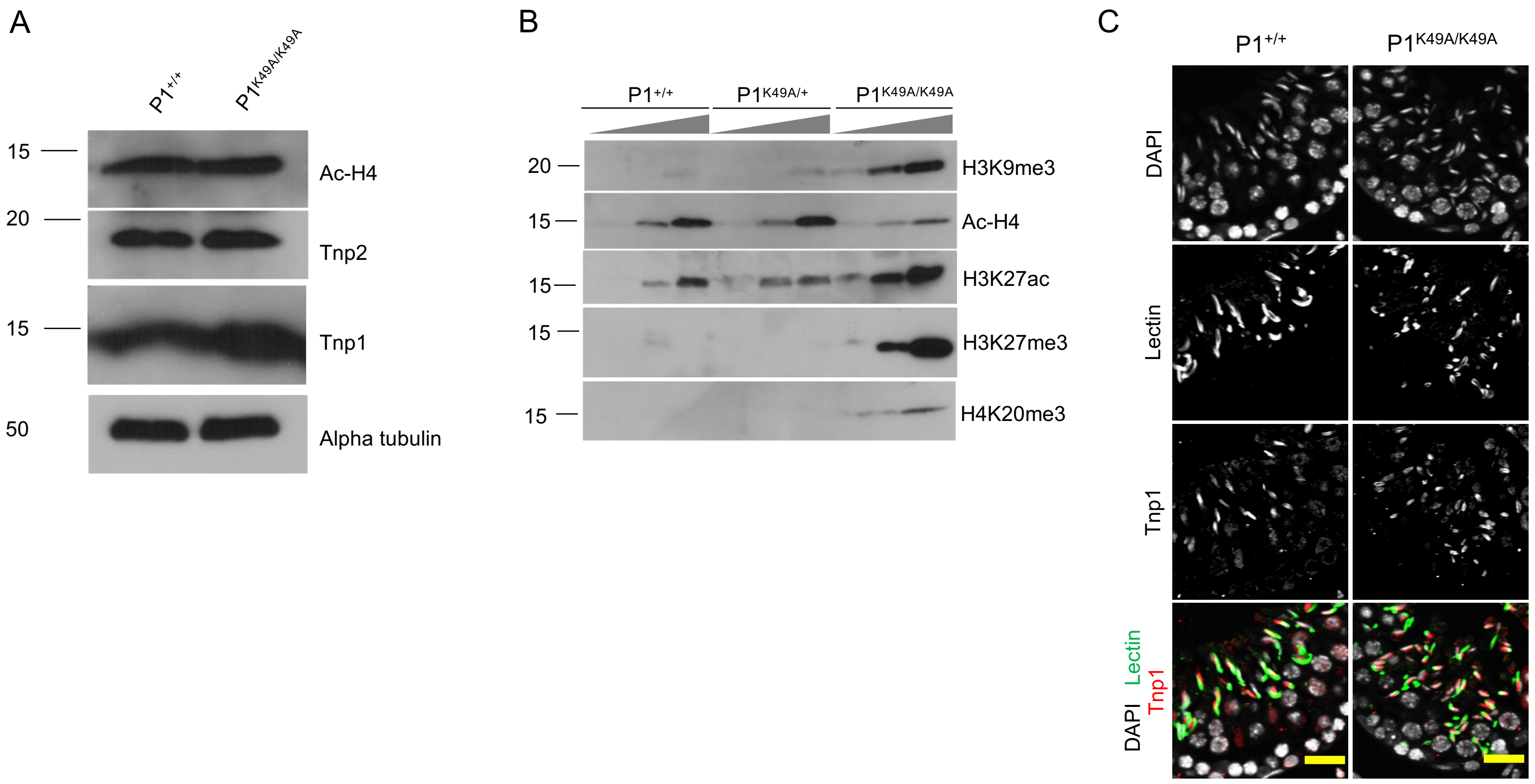


**Supplemental Figure 3: P1 K49A substitution alters sperm chromatin composition. (A)** Immunoblots of protein lysates from P1$^{+/+}$ and P1$^{K49A/K49A}$ elongating spermatid-enriched testes lysate illustrates no difference in ac-H4, TNP2, or TNP1 levels. **(B)** Immunoblotting of sperm protein extracts reveals an abnormal retention of modified histones in P1$^{K49A/K49A}$ sperm. Blots were loaded by total input sperm number. Exact sperm numbers for each antibody are provided in the Methods section. **(C)** Immunofluorescence staining of adult P1$^{+/+}$ or P1$^{K49A/K49A}$ testes cross sections stained for TNP1. Scale bars: 20 μm.

### Supplemental Figure 4

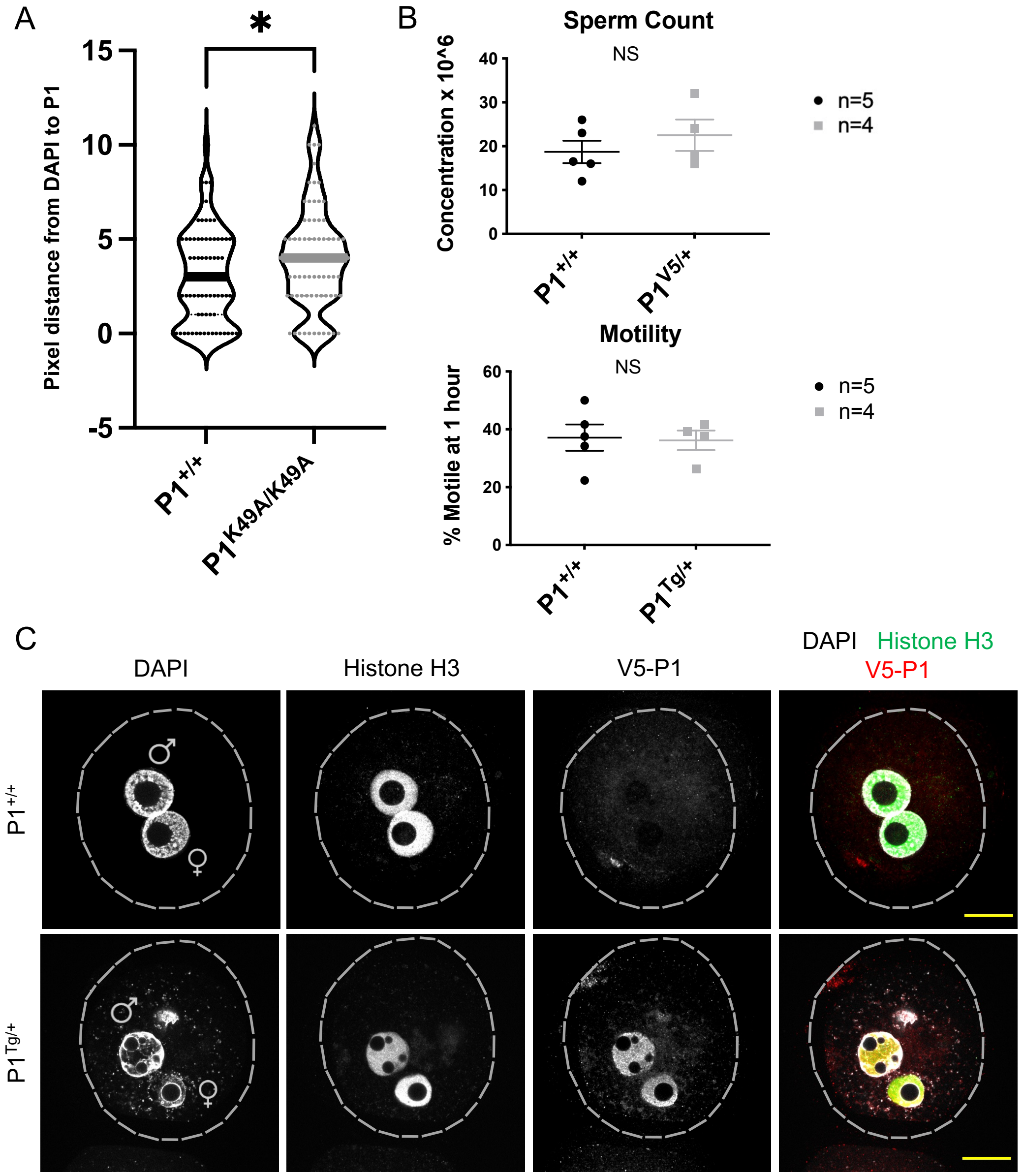


**Supplemental Figure 4:P1 K49A substitution results in decreased blastocyst formation and accelerated P1 dismissal from paternal chromatin . (A)** Pixel distance between edge of DAPI signal and highest P1 signal for $P1^{+/+}$ and $P1^{K49A/K49A}$ zygotes. A total of n=8 measurements were made for each embryo and a total of n=10 $P1^{+/+}$ zygotes and a total of n=12 $P1^{K49A/K49A}$ zygotes were used for quantification. Statistical test was performed using an unpaired t test, p=0.0123. **(B)** Sperm count and sperm motility for $P1^{+/+}$ and V5-$P1^{tg/+}$ males illustrate no difference upon addition of V5 tag. A total of n=5 $P1^{+/+}$ and n=4 V5-$P1^{tg/+}$ males were used. **(C)** Immunofluorescence of zygotes derived from $P1^{+/+}$ or V5-$P1^{tg/+}$ sperm, collected at 4hpf, and stained for V5 highlights localization of P1 to the maternal pronucleus. Scale bars: 20 µm.

### Supplemental Figure 5

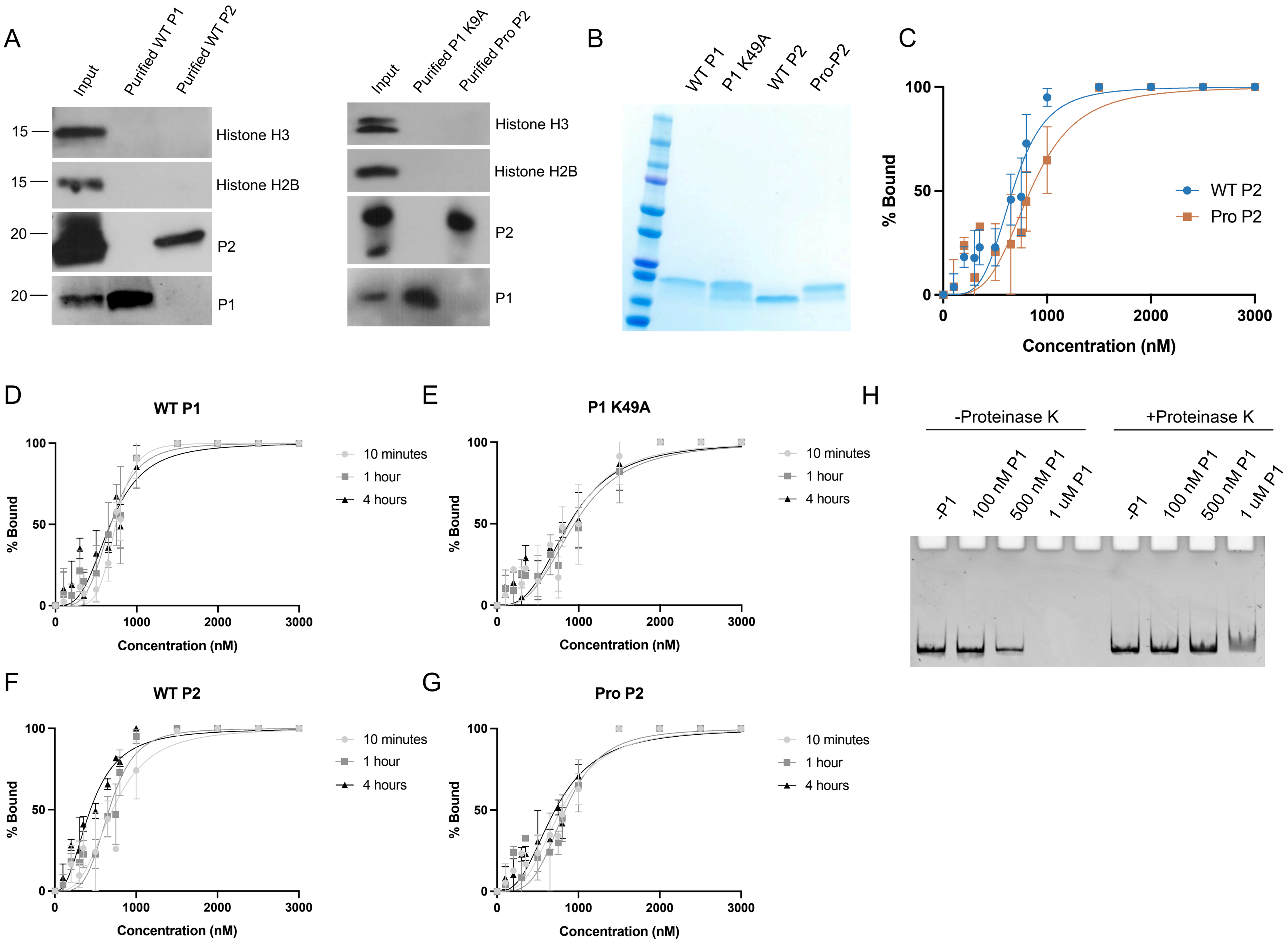


**Supplemental Figure 5: P1 K49A substitution negatively affects DNA binding. (A)** Immunoblot of input (prior to gel filtration chromatography) protein, purified P1 (WT or K49A), and purified P2 (WT or pro P2) illustrating efficient separation of P1 and P2 from each other, as well as the absence of histones in the final purified protein. **(B)** Coomassie-stained SDS-PAGE gel of purified proteins illustrating high purity and equivalent concentrations. **(C)** Quantification of the binding affinities of WT P2 and pro P2 to a linear ~300 bp DNA fragment. $K_{d,app}$ values were calculated using the Hill equation and were taken from at least 3 technical replicates per protein. **(D-G)** Quantification of binding affinities of WT P1 (D), P1 K49A (E), WT P2 (F), and pro P2 (G) after 10 minutes, 1 hour, or 4 hours of equilibration with DNA. Data were fit to a Hill curve and were taken from at least 3 technical replicates per protein. **(H)** Proteinase K treatment of EMSA reactions after 1 hour of equilibration with DNA.

### Supplemental Figure 6

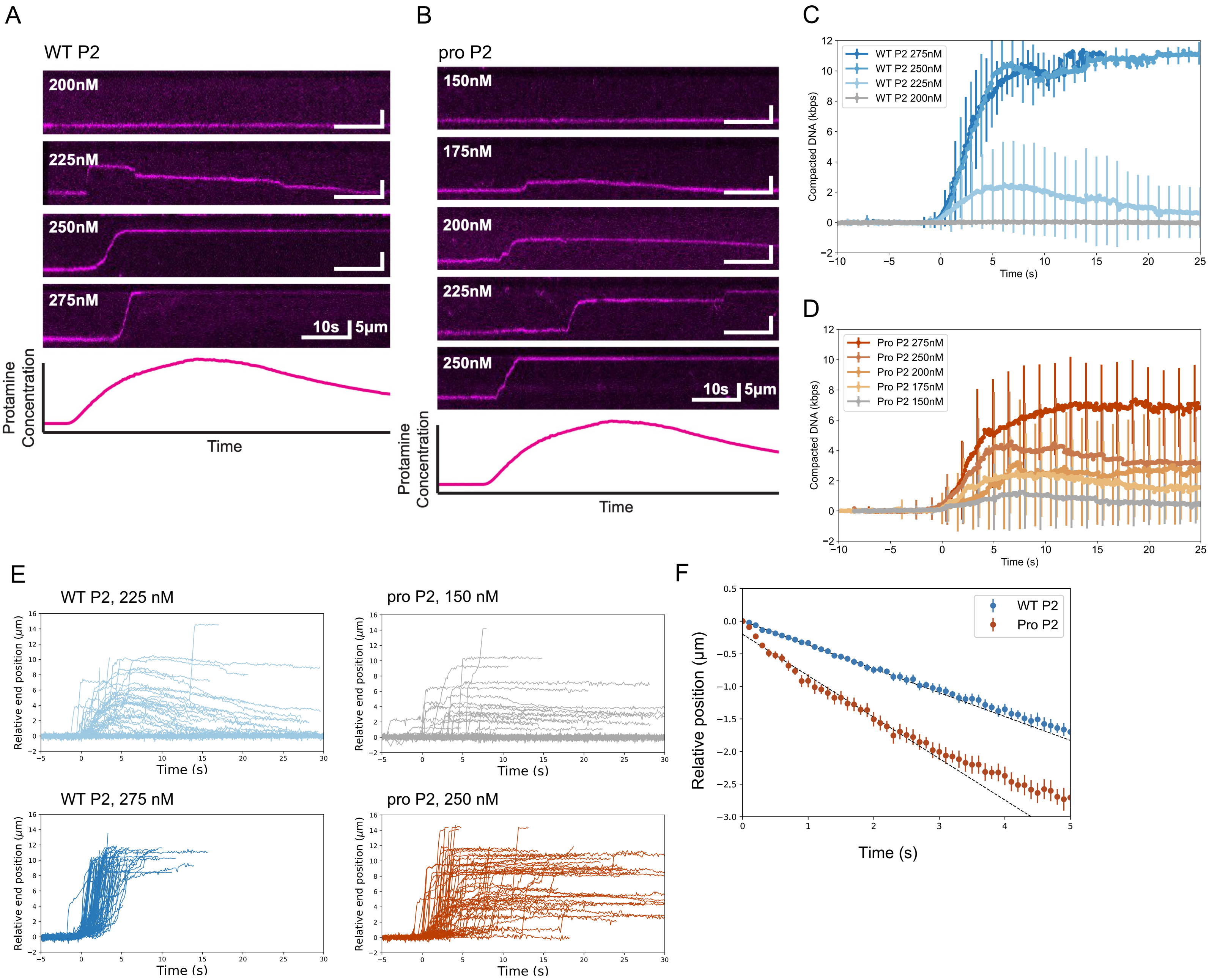


**Supplemental Figure 6: P1 K49A substitution alters DNA compaction and decompaction kinetics *in vitro*. (A)** Representative kymographs of WT P2 induced DNA compaction at increasing protein concentrations. **(B)** Representative kymographs of pro P2 induced DNA compaction at increasing protein concentrations. **(C)** Average DNA compaction by WT P2 at increasing concentrations. Error bars represent standard deviations (n=71 traces for 200 nM, n=63 for 225 nM, n=95 for 250 nM, and n=108 for 275 nM). **(D)** Average DNA compaction by Pro P2 at increasing concentrations. Error bars represent standard deviations (n=74 traces for 150 nM, n=54 for 175 nM, n=62 for 200 nM, n=64 for 225 nM, and n=65 for 250 nM). **(E)** Traces of individually tracked DNA molecules over time at low or high concentration of either WT P2 (left panels) or pro P2 (right panels) illustrating cooperative behavior. **(F)** Decompaction of DNA initially compacted by WT P2 and pro P2 over time illustrates differences in decompaction rates. Error bars represent SEM.
